# Enhancer Reprogramming Confers Dependence on Glycolysis and IGF signaling in KMT2D Mutant Melanoma

**DOI:** 10.1101/507327

**Authors:** Mayinuer Maitituoheti, Emily Z. Keung, Ming Tang, Liang Yan, Hunain Alam, Guangchun Han, Ayush T. Raman, Christopher Terranova, Sharmistha Sarkar, Elias Orouji, Samir B. Amin, Sneha Sharma, Maura Williams, Neha S. Samant, Mayura Dhamdhere, Norman Zheng, Tara Shah, Amiksha Shah, Jacob B. Axelrad, Nazanin E. Anvar, Yu-Hsi Lin, Shan Jiang, Edward Q. Chang, Davis R. Ingram, Wei-Lien Wang, Alexander Lazar, Min Gyu Lee, Florian Muller, Linghua Wang, Haoqiang Ying, Kunal Rai

## Abstract

Epigenetic modifiers have emerged as important regulators of tumor progression. We identified histone methyltransferase KMT2D as a potent tumor-suppressor through an *in vivo* epigenome-focused pooled RNAi screen in melanoma. KMT2D harbors frequent somatic point mutations in multiple tumor types. How these events contribute to tumorigenesis and whether they impart therapeutic vulnerability are poorly understood. To address these questions, we generated a genetically engineered mouse model of melanoma based on conditional and melanocyte-specific deletion of KMT2D. We demonstrate KMT2D as a bona fide tumor suppressor which cooperates with activated BRAF. KMT2D-deficient tumors showed substantial reprogramming of key metabolic pathways including glycolysis. Glycolysis enzymes, intermediate metabolites and glucose consumption rate were aberrantly upregulated in KMT2D mutant cells. The pharmacological inhibition of glycolysis reduced proliferation and tumorigenesis preferentially in KMT2D mutant cells. Mechanistically, KMT2D loss caused drastic reduction of H3K4me1-marked active enhancer states. Loss of distal enhancer and subsequent reduction in expression of IGFBP5 activated IGF1R-AKT to increase glycolysis in KMT2D-deficient cells. We conclude that KMT2D loss promotes tumorigenesis by facilitating increased usage of glycolysis pathway for enhanced biomass needs via enhancer reprogramming. Our data imply that inhibition of glycolysis or IGFR pathway could be a potential therapeutic strategy in KMT2D mutant tumors.

## INTRODUCTION

An important new theme that has emerged from the cancer genome sequencing studies in the past decade is genetic alterations in epigenetic regulators implicating epigenome as an important player in cancer progression (1,2). Loss-of-function missense and nonsense point mutations are observed to be highly prevalent across multiple tumor types in two families of chromatin regulators: 1) Histone H3K4 methyltransferase members including KMT2C and KMT2D; and 2) SWI/SNF complex members including SMARCA4, ARID1A, and PBRM1 (3). Although recent studies have begun to shed lights on roles of these proteins in cancer progression (4–7), we still have limited knowledge of why mutations in these proteins are selected over course of tumor progression.

We focus our studies on metastatic melanoma which is an aggressive cancer with a 5-year survival of less than 20% (8). In the past decade, the number of people affected by the disease have increased tremendously (8). Although the landscape of available treatment options has expanded for this disease in the form of immune checkpoint blockade agents and targeted agents (such as BRAFi and MEKi) (9), durable responses are observed in only a subset of patients leading to the death of several thousand people of this disease every year. Hence, other treatment strategies need to be further explored.

In cutaneous melanoma, mutations in epigenetic regulators, including IDH1/2, EZH2, ARID1A/1B, ARID2 and SMARCA4 have been observed at statistically significant frequencies (10,11). However, we have limited understanding of how specific mutant epigenetic proteins impact melanomagenesis. Functional studies have implicated the involvement of other epigenetic factors such as JARID1B (12), SETDB1 (13), TET2 (14) and histone variants (15,16) in melanoma progression. Systematic functional approaches are needed to elucidate how misregulation of epigenetic regulators impacts chromatin states and downstream gene expression programs during various stages of tumorigenesis. Detailed mechanistic understanding of melanomagenesis and the role of epigenetic regulators will also inform novel therapeutic strategies for patients whose tumors bear these mutations. We isolated KMT2D as the top hit in an *in vivo* RNAi screen focused on identification of epigenetic regulators that play tumor-suppressive function in melanoma. KMT2D is a histone H3 lysine 4 (H3K4) methyltransferase that primarily performs monomethylation, H3K4me1, which has been shown to be a marker of enhancer elements (17–20). KMT2D not only marks nucleosome with H3K4me1 but also recruits CBP/p300 that in turn acetylate these nucleosomes and hence lead to activation of these enhancers (18–20). Several studies have implicated enhancer aberrations as a hallmark of multiple tumor types including melanoma (21–31). However, most of them have focused on aberrant enhancer activation and little is known about how enhancer inactivation, which may result from loss of KMT2C/KMT2D function, influences tumor progression. We establish that KMT2D-deficient tumors may exhibit reduced enhancer activity that lead to the deregulation of energy metabolism pathways including glycolysis, thus providing a strategy for targeting of KMT2D-mutant cancers.

## RESULTS

### Identification of 8 Potential Tumor Suppressors including KMT2D through an RNAi Screen

We performed an RNAi screen (**Figure 1A**) to identify tumor-suppressor epigenetic regulators in melanoma. We used a well-characterized system of TERT-immortalized human primary foreskin melanocytes that harbor stably integrated dominant negative p53, CDK4^R24C^ and BRAF^V600E^ (32),(33) (passage n <15). These are referred as HMEL-BRAF^V600E^. When injected in NUDE mice, HMEL-BRAF^V600E^ cells forms visible tumors only after 22-24 weeks and with low penetrance (~10%-20%) (**Figures 1B**). In addition, this line is poised to switch to the tumorigenic state upon additional cooperative driver alterations such as PTEN loss (33) (**Figure 1B**). Hence, it is a good cell-based system for discovering tumor-promoting events as it provides a minimal yet sensitized tumorigenic background to identify moderate-to-potent tumor-suppressors. We also utilized this system for discovery of pro-tumorigenic epigenomic changes in melanoma (33).

**Figure 1:**
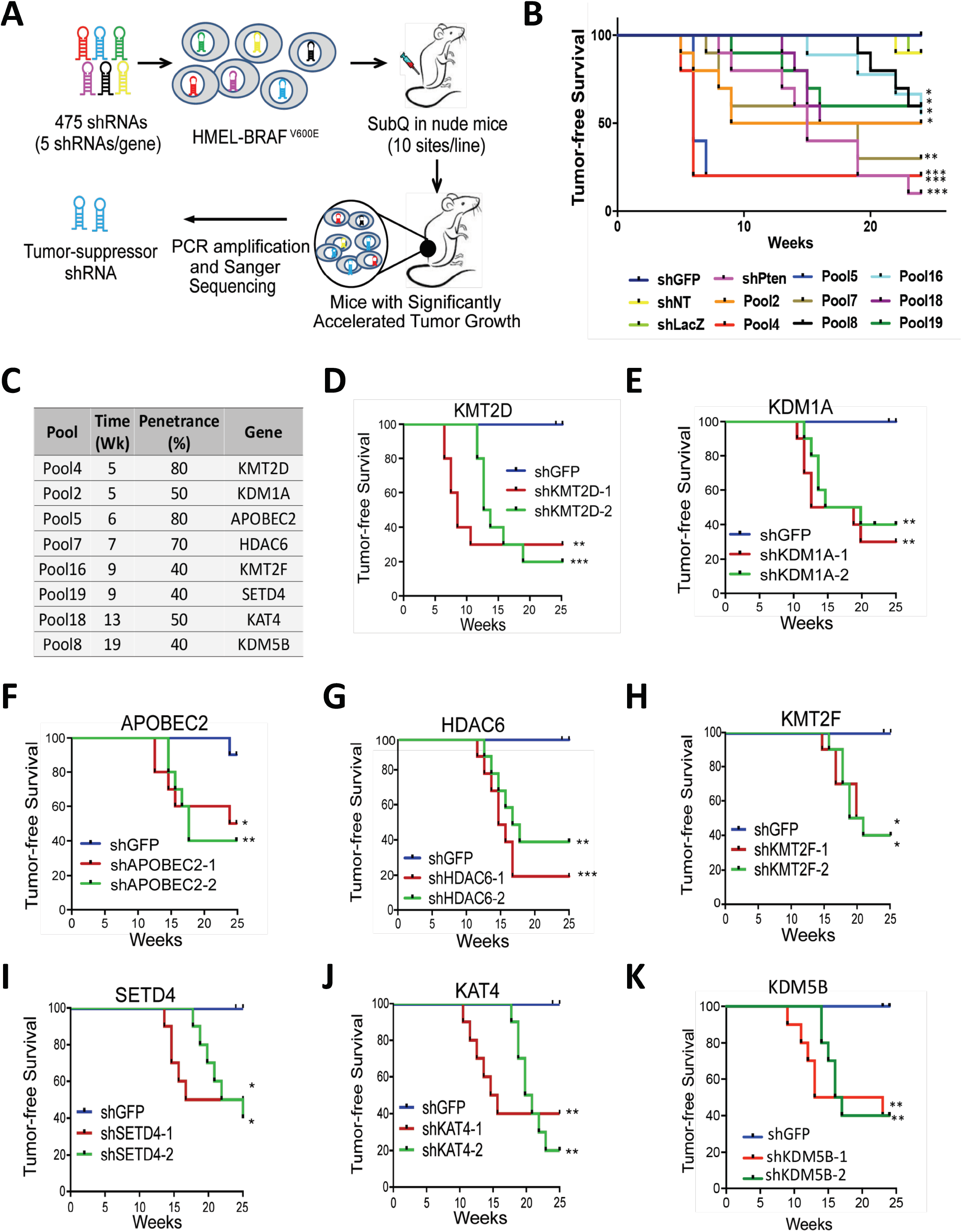
RNAi screen identifies potential melanoma tumor suppressor genes. **A.** Schematic of RNAi screen targeting epigenetic regulators to identify novel tumor suppressors in melanoma. HMEL-BRAF^V600E^ cells were transfected in a pooled fashion with 475 shRNAs targeting 95 epigenetic regulators (5 shRNAs/gene) in 19 experimental pools (25 shRNAs targeting 5 genes per experimental pool). Cells were orthotopically injected intradermally into the flanks of NCR-NUDE mice. Tumors that arose before the controls (shNT, shLuc and shGFP) were sequenced to identify the shRNA sequence. **B.** Kaplan-Meier curve showing tumor-free survival of mouse cohorts orthotopically injected with 1 million HMEL-BRAF^v600E^ cells transfected with pooled shRNAs from primary screen. Nineteen experimental pools (P1-P19), 3 negative control pools (shLuc, shGFP and shNT), and one positive control (shPTEN) were injected in 10 mice each and tumor formation was monitored over 25 weeks. Mantel-cox test *p < 0.05, **p < 0.01, and ***p<0.001; n = 10 per pool. Only pools that show significant acceleration are shown here. For data on not significant pools, please See Figure S1A. **C.** List of genes identified from their pools, the week of first appearance of tumor in the pool and the percentage of mice in respective cohort demonstrating accelerated tumorigenesis over control mice (penetrance). **D-N**, Kaplan-Meier curves showing tumor-free survival of mouse cohorts orthotopically injected with HMEL-BRAF^V600E^ cells stably expressing shRNAs against KMT2D (**D**), KDM1A (**E**), APOBEC2 (**F**), HDAC6 (**G**), KMT2F (**H**), SETD4 (**I**), KAT4 (**J**) and KDM5B (**K**). Mantel-cox test *p < 0.05, **p < 0.01, and ***p<0.001; n = 10 per arm.

In the current study, we constructed a shRNA expression vector library that included 475 shRNAs targeting 95 proteins known to regulate epigenetic processes including chromatin modification and nucleosome remodeling (**Table S1**). The HMEL-BRAF^V600E^ cells were transfected with 23 pools of shRNAs individually. Hereafter, “pool” refers to stably transfected HMEL-BRAF^v600E^ cells. Of these, 19 experimental pools contained 25 shRNAs each (5 shRNAs each for 5 genes selected randomly). The three negative pools contained one negative control shRNA each [shGFP, shLacZ and shNT (non-targeting)] and final pool harbored PTEN shRNA (shPTEN) as a positive control (**Figure 1B**). Briefly, 1 million cells were orthotopically injected intradermally in NUDE mice (10 sites) which were monitored for visible tumor formation over the subsequent 25 weeks (**Figures 1B, S1A**). Mice injected with cells from 8 of the 19 pools and the positive control (shPTEN) displayed significant acceleration of tumor formation compared to the negative controls. The first occurrence of tumor formation was at 5 weeks while tumor formation did not occur until 22 weeks in the negative control mice and multiple pools (**Figures 1B-C, S1A**).

We next identified the shRNAs enriched in tumors harvested from that exhibited significantly accelerated tumor formation by isolating tumor DNA and performing Sanger sequencing of the pLKO amplified region containing shRNA (list of genes in **Figure 1C**). We identified 8 unique shRNAs each from 8 pools that significantly accelerated tumor formation. To validate the results of the screen, we knocked down each of the 8 candidate genes individually using at least two independently validated shRNAs in HMEL-BRAF^V600E^ and widely used WM115 (BRAF^V600E^ mutant) melanoma cells (**Figures S1B-I**) and tested tumor formation efficiency (**Figures 1D-K and S1J-P**). All 8 genes (KMT2D, KDM1A, APOBEC2, HDAC6, KMT2F, SETD4, KAT4 and KDM5B) were validated as tumor suppressor candidates as knockdown of these genes in both HMEL-BRAF^V600E^ and WM115 cells resulted in accelerated tumor formation (p < 0.05) (**Figures 1D-K and S1J-P**). In addition, knockdown of a subset of these genes in HMEL-BRAF^V600E^ cells also promoted invasion *in vitro* in a Boyden chamber assay (**Figure S1Q**). KMT2D was the most potent hit as mice injected with cells with stable KMT2D knockdown developed tumor appearance at the earliest interval and with the highest penetrance compared to negative controls (**Figures 1C-D and 2A**). Among the rest, KDM5B has been previously implicated in melanomagenesis where it is believed to control the maintenance of melanoma stem cells(12,34).

**Figure 2:**
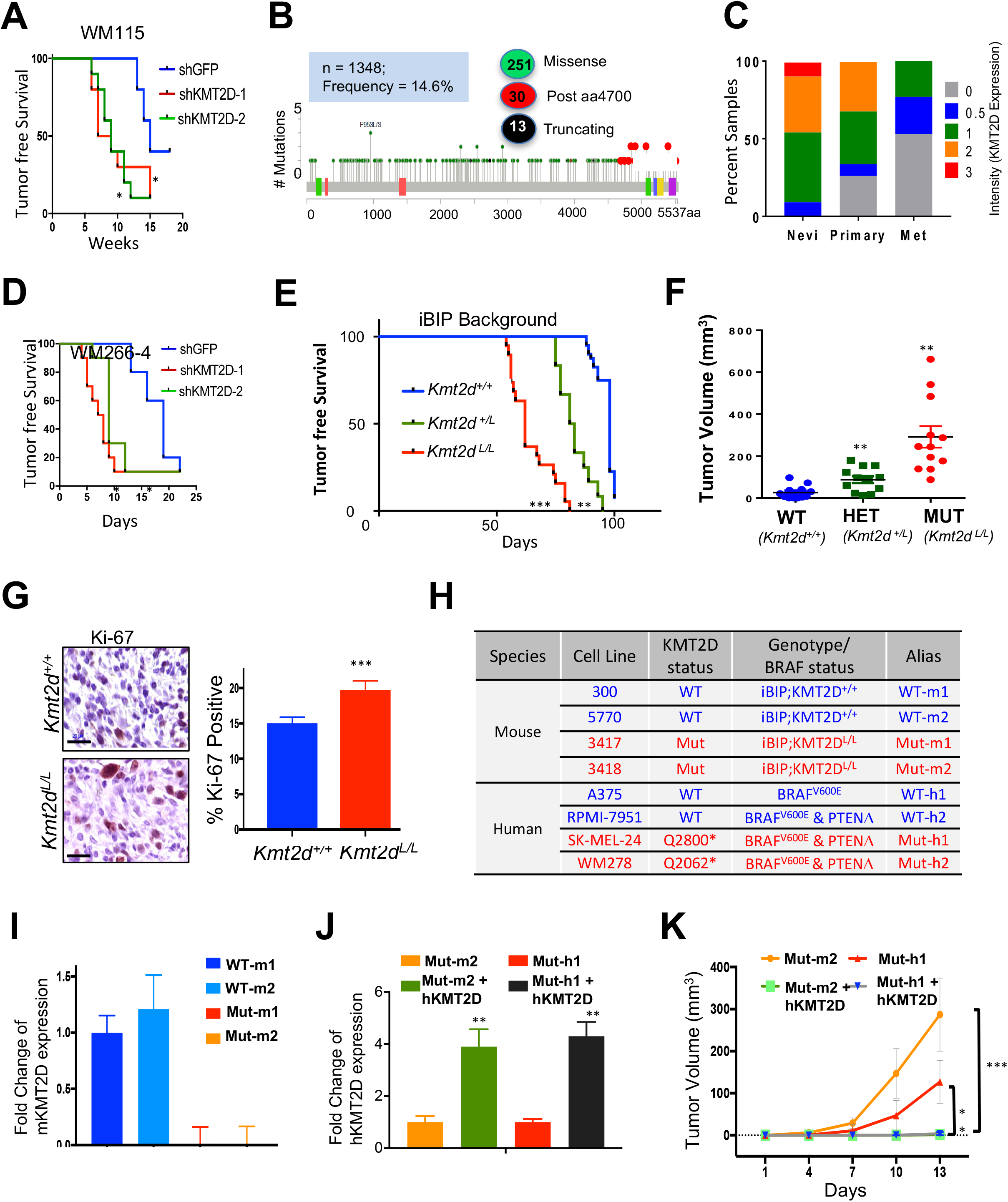
KMT2D functions as a tumor-suppressor in BRAF^V600E^ mutant melanomas. **A, D.** Kaplan-Meier curve showing tumor-free survival of mouse cohorts orthotopically injected with WM115 (**A**) and WM266-4 (**D**) cells stably expressing shRNAs against KMT2D. Mantel-cox test *p < 0.05. **B.** Schematic of KMT2D protein showing missense mutations seen across all melanoma studies deposited in cBio portal. Green filled circles indicate missense mutations, black filled circles indicate truncating mutations and red circles indicate functional mutations occurring after amino acid residue 4700. Colored boxes within KMT2D schematic show different protein domains. **C.** Stacked bar chart showing percent of nevi (n = 18), primary melanoma (n = 62) and metastatic melanoma (n = 22, labelled ‘Met’) samples with the various intensity (of KMT2D expression) categories (0, 0.5, 1, 2, 3 as shown in the legend). The p-value for difference in average intensity of KMT2D expression between primary and nevi was < 0.05. Similarly, p-value for difference in average intensity of KMT2D expression between primary and metastatic melanoma was < 0.05. **E.** Kaplan-Meier curve of auricular tumor-free survival in KMT2D WT (KMT2D+^/^+, blue), KMT2D heterozygous (KMT2D^L/+^, green) and KMT2D mutant (KMT2D^L/L^, red) mice in an iBIP (*Tyr-Cre^ERT2^, Rosa26-rtta, TetO-BRAF^V600E^, PTEN^L/L^, INK/ARF^L/L^*) background that were treated with doxycycline (2mg/ml, *ad libitum*) and 4-OHT (1μM, topical). X-axis refers to days after 4-OHT application. Mantel-cox test *p < 0.05, **p < 0.01, and ***p<0.001. **F.** Tumor burden of KMT2D WT (KMT2D^+/+^, blue), KMT2D heterozygous (KMT2D^L/+^, green) and KMT2D mutant (KMT2D^L/L^, red) mice in iBIP background at 89 days post 4-OHT application. **G.** Images of Ki-67 stained (standard Immunohistochemistry) (40X) melanoma tumors from *iBIP;KMT2D^+/+^* and *iBIP;KMT2D^L/L^* mice. Right panel shows percent of Ki-67 stained cells across 5 different fields of 100 cells each. **H.** Table showing the human melanoma lines and *iBIP;KMT2D* mouse model-derived cells that were used in the functional studies through the rest of the figures. **I.** Bar graph showing KMT2D expression levels in WT-m1, WT-m2, Mut-m1 and Mut-m2 to demonstrate loss of KMT2D mRNA. Y-axis represents fold change of the gene expression compared to 28S and normalized to average values in WT-m1. See Figure S2i for protein levels. **J.** Bar graph showing KMT2D expression levels in Mut-m2 and Mut-h1 lines that express doxycycline-inducible full length KMT2D protein. Y-axis represents fold change of the gene expression compared to 28S and normalized to Mut-m2 or Mut-h1 lines. See Figure S2i for protein levels. **K.** Graph showing tumor volume of NUDE mice injected with KMT2D mutant murine (Mut-m2) or human (Mut-h1) cells harboring inducible KMT2D expression vector that leads to KMT2D overexpression upon application of doxycycline. Top 5 GO terms for upregulated genes (FDR < 0.05, FC >2) between KMT2D mutant murine cells and KMT2D wild type cells by total RNA-Seq analysis.

### GEMM Model Confirms KMT2D is a Potent Tumor Suppressor in Melanoma

We searched published melanoma genomic studies to identify patients whose tumors harbor genetic aberrations in the potential tumor suppressor genes discovered through the screen. We observed that ~15% of melanoma cases identified harbored missense mutations in KMT2D whereas ~5%-8% of patients harbored missense mutations in KAT4 (**Figures 2B and S2A**) (35). As KMT2D mutations are prevalent (36–52) and this gene is increasingly reported to be a potential tumor suppressor across other tumor types (4–7,36,53), we next sought to deeply characterize the mechanism of action of KMT2D in melanoma, particularly as the strongest phenotype in RNAi screen was seen with the KMT2D loss. A subset of the missense mutations in KMT2D were truncating or frameshift insertions/deletions (4.4%) that likely abrogates histone methyltransferase activity (**Figure 2B**). In addition, 10% of all missense mutations occurred distal to amino acid residue 4700 which were shown to disrupt histone methyltransferase activity in a previous study (7). Together, we categorize these set of mutations – truncating, frameshift and post4700aa - as ‘functional’ driver mutations for KMT2D. Although we make use of this stringent criteria as a deterministic measure for KMT2D-deficient tumors so that we can delineate its mechanism of action, it is not a reflection of all KMT2D somatic variants that may produce a nonfunctional KMT2D protein.

First, we checked if, in addition to mutations, KMT2D mRNA and protein levels were also altered in human melanoma. Staining of a tissue microarray harboring 100 cases of nevi, primary melanomas and metastatic melanomas showed significant progressive loss of KMT2D protein levels in primary and metastatic melanomas (**Figure 2C**). Similar trend was also observed in mRNA expression of KMT2D as identified by assessment of publically available melanoma progression transcriptomic datasets (54,55) (**Figure S2B**) suggesting KMT2D regulation at both the level of gene expression and somatic mutations.

Functionally, knockdown of KMT2D by two different shRNAs led to increased tumor burden in two clonal variants of HMEL-BRAF^V600E^, WM115 and WM266-4 cells (**Figures 1D, 2A, 2D and S2C-G**). In addition, an increase in soft agar colony formation (**Figure S2H**) and invasion (**Figure S2I**) was observed *in vitro*. To further verify the role of KMT2D in melanoma in a specific genetic context, we utilized a conditional mouse model of KMT2D that harbors Lox sites flanking exons 16-20, thereby leading to deletion of this gene in tissue-specific manner(4). Melanocyte-specific deletion with Tyr–Cre^ERT2^ did not result in the formation of melanomas (data not shown) and thus these mice were crossed with a previously published doxycycline- and a tamoxifen-inducible mouse model of BRAF^V600E^ melanoma (*iBIP = Tyr-Cre^ERT2^, Rosa26-rtta, TetO-BRAF^V600E^, PTEN^L/L^, INK/ARF^L/L^*). Tamoxifen application on the ears of KMT2D mutant iBIP mice resulted in a drastic acceleration of tumorigenesis in comparison to KMT2D wild-type (WT) iBIP mice (**Figure 2E-F**). Intriguingly, the heterozygous mice also showed significant acceleration of auricular tumor burden (**Figure 2E-F**). Further, increased proliferation and melanocyte origin was confirmed by immunohistochemical analysis (IHC) for Ki-67 and tyrosinase, respectively (**Figures 2G, S2J**).

Next, we derived cell lines from the tumors of 2 KMT2D wild-type (WT, iBIP-KMT2D^+/+^) and 2 mutant (Mut, iBIP-KMT2D^L/L^) models (**Figure 2H**) and confirmed for the genotype of all alleles. The 2 KMT2D WT iBIP cell lines (labeled as WT-m1 and WT-m2 for WT mouse 1 and 2) and 2 KMT2D mutant cell lines (labeled as Mut-m1 and Mut-m2 for mutant mouse 1 and 2) (**Figure 2H**). These lines were verified for loss of KMT2D mRNA by qPCR (**Figure 2I**) and protein by immunofluorescence (**Figure S2K**). The phenotypes observed in KMT2D mutant lines were dependent on the loss of this gene as overexpression of full-length KMT2D (**Figures 2J, S2K**) reduced tumor burden *in vivo* (**Figure 2K**) in immunodeficient NUDE mice. To assess relevance in humans, we performed all of the follow-up experiments in 2 KMT2D WT human melanoma lines (A375 and RPMI-7951 which are referred to hereafter as WT-h1 and WT-h2) and 2 KMT2D mutant human melanoma lines (SKMEL-24 and WM278 which are referred to hereafter as Mut-h1 and Mut-h2) (**Figure 2H, S2K**). SKMEL-24 (Mut-h1) and WM278 (Mut-h2) harbor truncating mutations at Q2800 and Q2062 respectively(56).

### Hyperactive Glycolysis in KMT2D Mutant Melanomas is a Targetable Pathway

To determine the molecular phenotype conferred by KMT2D loss, we performed an RNA-Seq-based transcriptome profiling experiment in the KMT2D WT and mutant murine melanoma lines. We identified 1761 genes that were uniquely overexpressed in the KMT2D mutant compared to WT conditions (FDR < 0.05, FC >2, n = 3) and 1443 which were repressed. Genes overexpressed in KMT2D mutant cells were enriched for pathways related to immune response, cell adhesion, epithelial-to-mesenchymal transition as well as various metabolic pathways including the “hexose metabolic pathway” or glycolysis (**Figure 3A-B, S3A-B**). Similar pathways, including Glycolysis, were also found to be upregulated in KMT2D mutant human melanomas upon analyses of primary melanoma tumors from a published TCGA study (**Figures 3C, S3C**). A survey of pan-cancer TCGA data suggested that energy metabolism pathways including glycolysis were activated across 6 other tumor types (BLCA, urothelial bladder carcinoma; CESC, squamous cell carcinoma and endocervical adenocarcinoma; HNSC, head and neck squamous cell carcinoma; LUSC, lung squamous cell carcinoma; UCEC, uterine corpus endometrial carcinoma; STAD, stomach adenocarcinoma) that harbor ‘functional’ KMT2D driver mutations (**Figures 3D, S3D and Table S2**). We observed drastic upregulation of 10 of 12 glycolysis pathway enzyme genes (GLUT1, HK1, GPI1, PFKA, ALDOC, TPI1, GAPDH, PGK1, PGAM1 and ENO1) by qPCR in KMT2D mutant lines in comparison to WT lines in both human and murine models (**Figures 3E, 3F, S3E**). Similarly, the rescue of mutant lines with full-length WT KMT2D reduced their expression (**Figure 3G**). Higher expression of ENO1, PGK1 and PGAM1 was confirmed in KMT2D mutant iBIP melanoma tumors by immunohistochemical analysis (**Figure 3H**). Quantitation of glucose uptake and lactate production confirmed upregulation of glycolysis in the KMT2D mutant lines (**Figures 3I-J**) which was reduced upon KMT2D overexpression (**Figure 3K**). In addition, mass-spectrometry-based quantitative measurement of glycolysis intermediate metabolites showed higher levels of fructose-1,6-biphosphate, D-glyceraldehyde-3-phosphate, 1,3-diphosphateglycerate and pyruvate in the KMT2D mutant murine line (**Figure 3L**). Consistent with this higher glycolysis rate in KMT2D mutants, they grew poorly in low glucose media compared to high glucose media which likely resulted from rapid exhaustion of glucose in the media (**Figure S3E-F**). A trivial explanation for increase in glycolysis in KMT2D mutant cells would be the higher proliferative potential of these cells compared to wild type cells. However, contradictory to this hypothesis, we observe that KMT2D mutant cells proliferate slower than wild type cells *in vitro* (**Figure S3E-F**) despite increased proliferation and tumorigenesis *in vivo*. Together, these data provide the evidence of activation of glycolysis in KMT2D mutant melanomas (**Figure 3M**).

**Figure 3:**
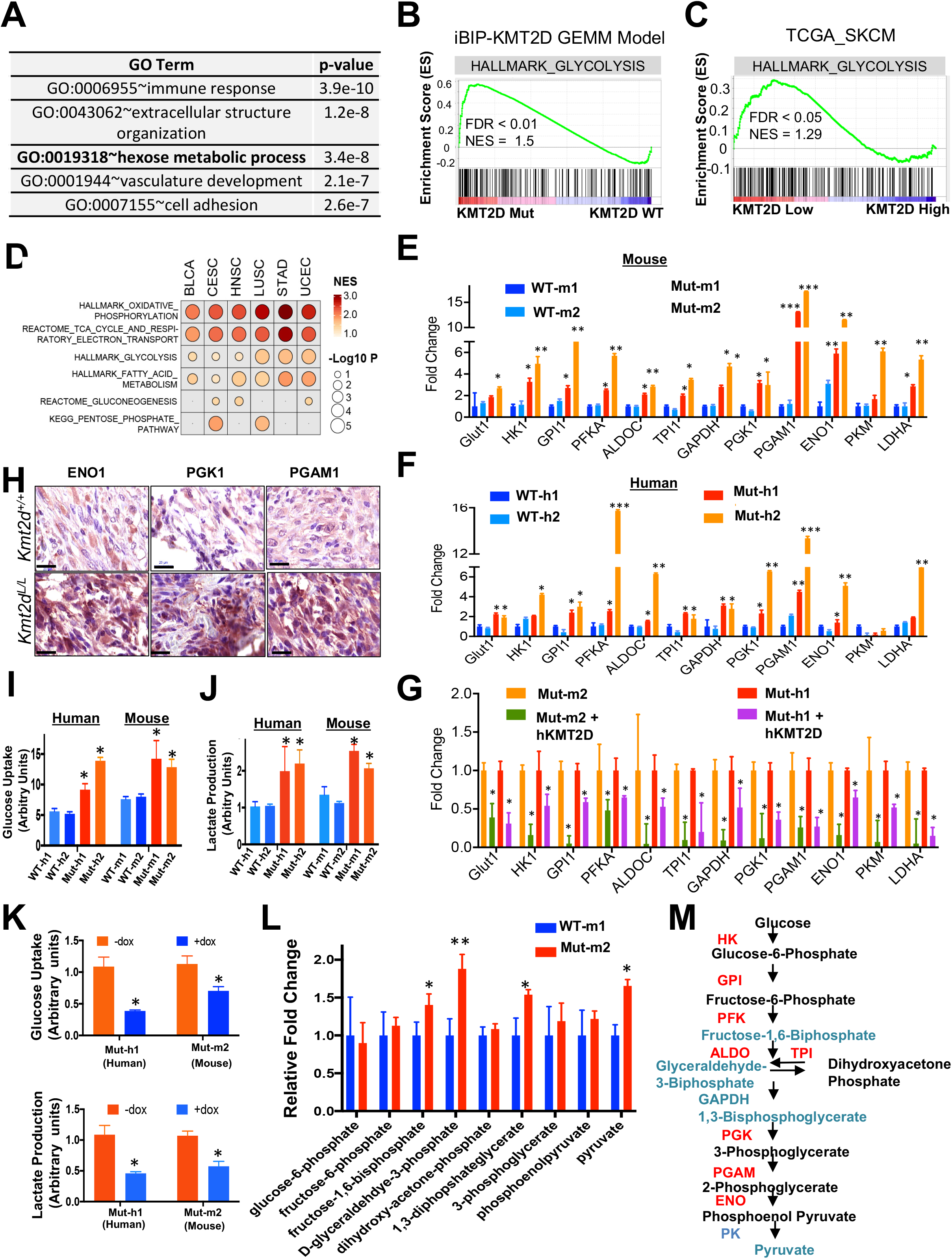
KMT2D mutated tissues exhibit aberrant activation of glycolysis. **A.** Top 5 GO terms for upregulated genes (FDR < 0.05, FC >2) between KMT2D mutant murine cells and KMT2D wild type cells by total RNA-Seq analysis. **B.** Enrichment plot for HALLMARK glycolysis pathway in all differentially expressed genes (FDR < 0.05, FC >2) between KMT2D mutant murine cells and KMT2D WT cells by total RNA-Seq analysis. Each black bar represents a gene in the pathway. **C.** GSEA plot of the HALLMARK glycolysis pathway in differentially expressed genes between human primary melanomas with low versus high KMT2D expression (n = 10 for each group) from the TCGA-SKCM cohort. Each black bar represents a gene in the pathway. **D.** GSEA of different MSigDB energy metabolism pathways in differentially expressed genes between KMT2D mutant (carrying truncation, frameshift and post4700aa missense) and WT human tumors from six TCGA tumor groups where KMT2D mutant tumors n > 10. **E-G.** Bar graph showing the relative expression pattern of 12 glycolysis enzyme genes (compared to 28S) in KMT2D mutant and WT murine (**E**) and human (**F**) cells (details of the system in Figure 2I) as well as in Mut-m1 and Mut-h1 cells with dox-inducible rescue of full length WT KMT2D expression (**G**). **H.** Immunohistochemistry images demonstrating expression of ENO1, PGK1 and PGAM1, encoded by three glycolysis genes, in *iBIP;KMT2D*^+/+^ and *iBIP;KMT2D^L/L^* melanoma tumors. **I-K.** Graph showing measurement of glucose uptake and lactate production in KMT2D mutant and WT murine (**I**) and human (**J**) cells (details of the system in Figure 2i) as well as in the KMT2D mutant mouse Mut-m1 and human Mut-h1 cells with doxycyclin-inducible rescue of full length WT KMT2D expression (**K**). **L.** Bar graph showing relative levels of various metabolite intermediates produced during glycolysis pathway as measured by selected reaction monitoring tandem mass spectrometry. Asterisk denotes *p < 0.05, ** p < 0.001. **M.** Schematic of the glycolysis pathway showing aberrantly activated glycolysis enzymes and (in red) and metabolites (in green) KMT2D mutant compared to in WT conditions.

Next, we tested whether the aberrantly activated glycolysis pathway contributed to the increased tumorigenic potential of KMT2D mutant melanomas. Inhibition of the glycolysis pathway using 3 different inhibitors - 2-DG (glucose competitor), Pomhex (a novel ENO1 inhibitor(57)) and Lonidamine (Hexokinase inhibitor) – selectively reduced proliferation of KMT2D mutant melanoma cells in comparison to KMT2D wild-type melanoma cells in both murine as well as human systems (**Figures 4A-D and S4A-B**). This effect was more pronounced in low glucose conditions compared to high glucose media (**Figure S4C-D**). This preferential effect of 2-DG on KMT2D mutant murine and human cell lines was rescued with the expression of WT KMT2D (**Figures 4E-F**). Consistent with the *in vitro* data, tumors formed by xenotransplantation of KMT2D mutant lines were more sensitive to 2-DG treatment in NUDE mice (**Figure 4G-H and S4E-H**). Importantly, we did not observe preferential growth inhibition of KMT2D mutant murine cells in comparison to WT cells by a OxPhos inhibitor, IACS-10759 (**Figure S4I**). Together, these data suggest that upregulated glycolysis is an important contributor to enhanced tumorigenesis in KMT2D mutant melanomas and suggest a potential therapeutic strategy in this genetic context.

**Figure 4:**
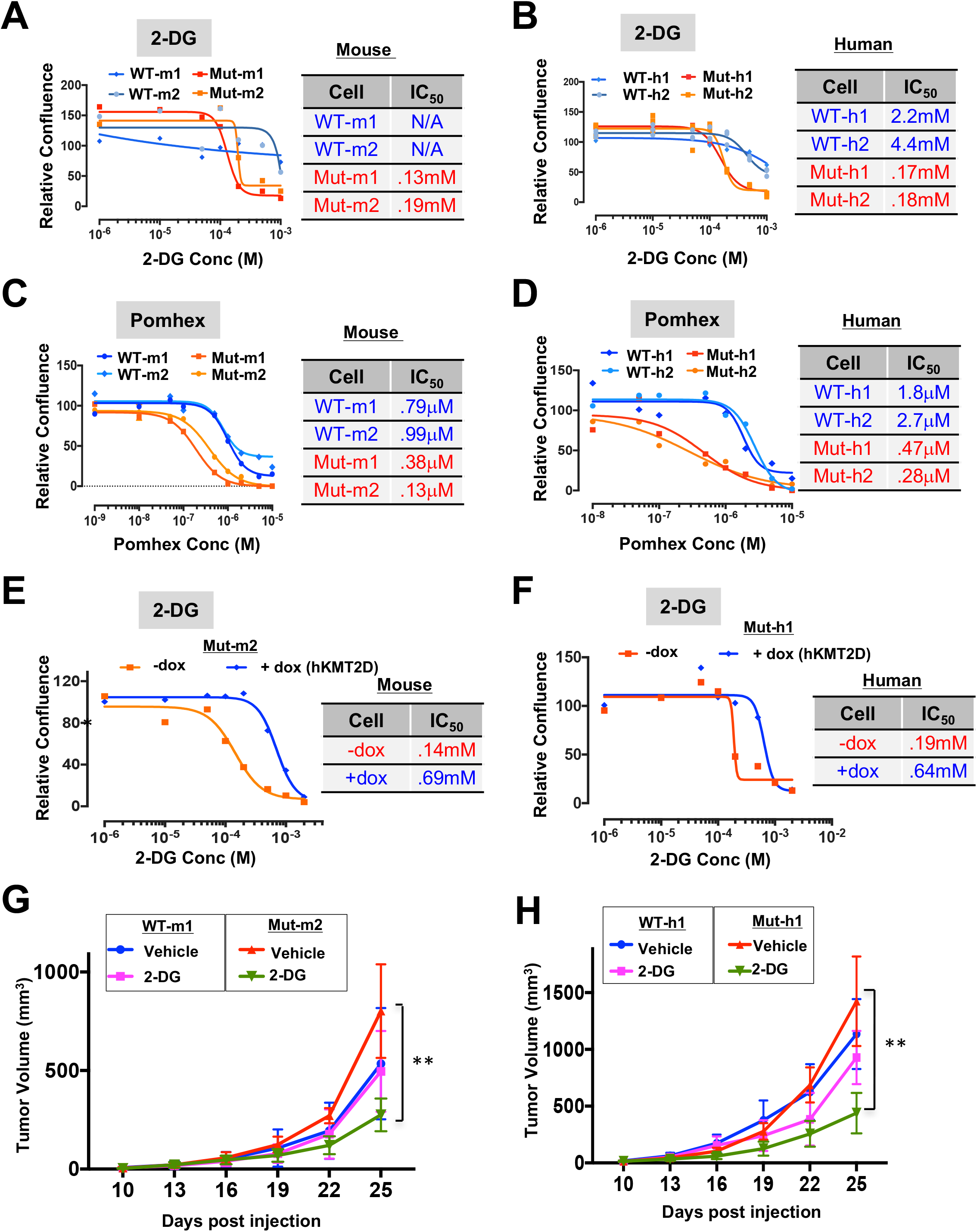
Inhibition of glycolysis preferentially impacts KMT2D mutant cells. **A-D,** Growth curves for KMT2D mutant and WT murine (**A, C**) and human (**B, D**) melanoma cells treated with varying concentrations of 2-Deoxy-D-Glucose (**A, B**) or Pomhex (**C, D**). Relative confluence at 96-hours post treatment are plotted and IC_50_ values are shown in the accompanying table. **E-F.** Growth curves for KMT2D mutant mouse Mut-m2 (**E**) and human Mut-h1 (**F**) melanoma cells that express inducible KMT2D (with 10mg/ml doxycycline application). Relative confluence at 96-hours post treatment are plotted and IC50 values are shown in the accompanying table. **G-H.** Line plot showing average tumor volumes for mice (n = 5 per group) injected with KMT2D mutant and WT murine (G) and human (H) melanoma cells and treated with 2-DG (500mg/kg) every other day.

### H3K4me1-marked Enhancer and Super-enhancer Reprogramming Occurs in KMT2D Mutant Melanoma

We examined total and genome-wide levels of H3K4 marks as KMT2D is known to harbor histone methyltransferase activity toward multiple H3K4 methylation states and impacts H3K27ac patterns (4,19,20,58,59). KMT2D mutant murine cells harbored lower levels of total H3K4me1 and H3K27ac marks in comparison to WT cells (**Figure 5A**). Consistently, H3K4me1 levels were elevated upon KMT2D reexpression (**Figure S5A**). Next, we determined chromatin states in murine melanoma KMT2D mutant and WT tumors using ChIP-seq for the histone modifications H3K4Me1 (enhancers), H3K4Me3 (promoters), H3K27Ac (active), H3K79Me2 (transcription) and H3K27Me3 (polycomb-repressed) (60) in line with studies from NIH Roadmap project(61). Chromatin state calls using a 10-state ChromHMM model representing various epigenomic states including promoters (States 3 and 4), enhancers (States 2, 5 and 6), polycomb-repressed (State 9), transcribed (State 1) and unmarked (States 8, and 10) (**Figure 5B**). Chromatin state transition between KMT2D mutant and WT cells identified state 6 (active enhancer, H3K4me1 high) to 10 (Low) as the most prominent transition which was associated with loss of H3K4me1 and H3K27ac based enhancers (**Figure 5C**). The other two prominent changes [i.e. State 2 (Transcribed enhancer) to 1 (Transcribed), and State 3 (Transcribed 5’ and 3’ promoter) to 2 (Transcribed enhancer)] were associated with the loss of H3K4me1 and H3K4me3 respectively (**Figure 5C**). An examination of average intensities of individual H3K4me1 and H3K27ac marks across the genome showed a significant reduction of H3K4me1 and little/no change in H3K27ac (**Figures 5D-E**). Similarly, we observed changes in H3K4me1-based superenhancer regions but not those called by H3K27ac signal (**Figure S5B-C**). Importantly, we also noticed pronounced increase in average intensities of H3K27me3 peaks in KMT2D mutants compared to WT samples on enhancer loci that lose H3K4me1 mark which could imply likely transcriptional repression of a subset of genes (**Figure 5F**). H3K27me3 peaks showed modest enrichment genome-wide as well (**Figure S5D**). This could be due to loss of function of H3K27me3-specific demethylase, KDM6A, which is known to be an obligate partner of KMT2D (59). On the contrary, we did not notice much change in H3K79me2 and H3K4me3 enrichment between KMT2D wild type and mutant tumors (**Figure S5E-F**). We also identified active enhancer loss associated with important melanoma associated pathways including immune pathways, apoptosis signaling pathway and p53 pathway by glucose deprivation (**Figure S5G**). These data suggest that KMT2D loss results in significant reprogramming of the enhancer landscape in melanoma.

**Figure 5:**
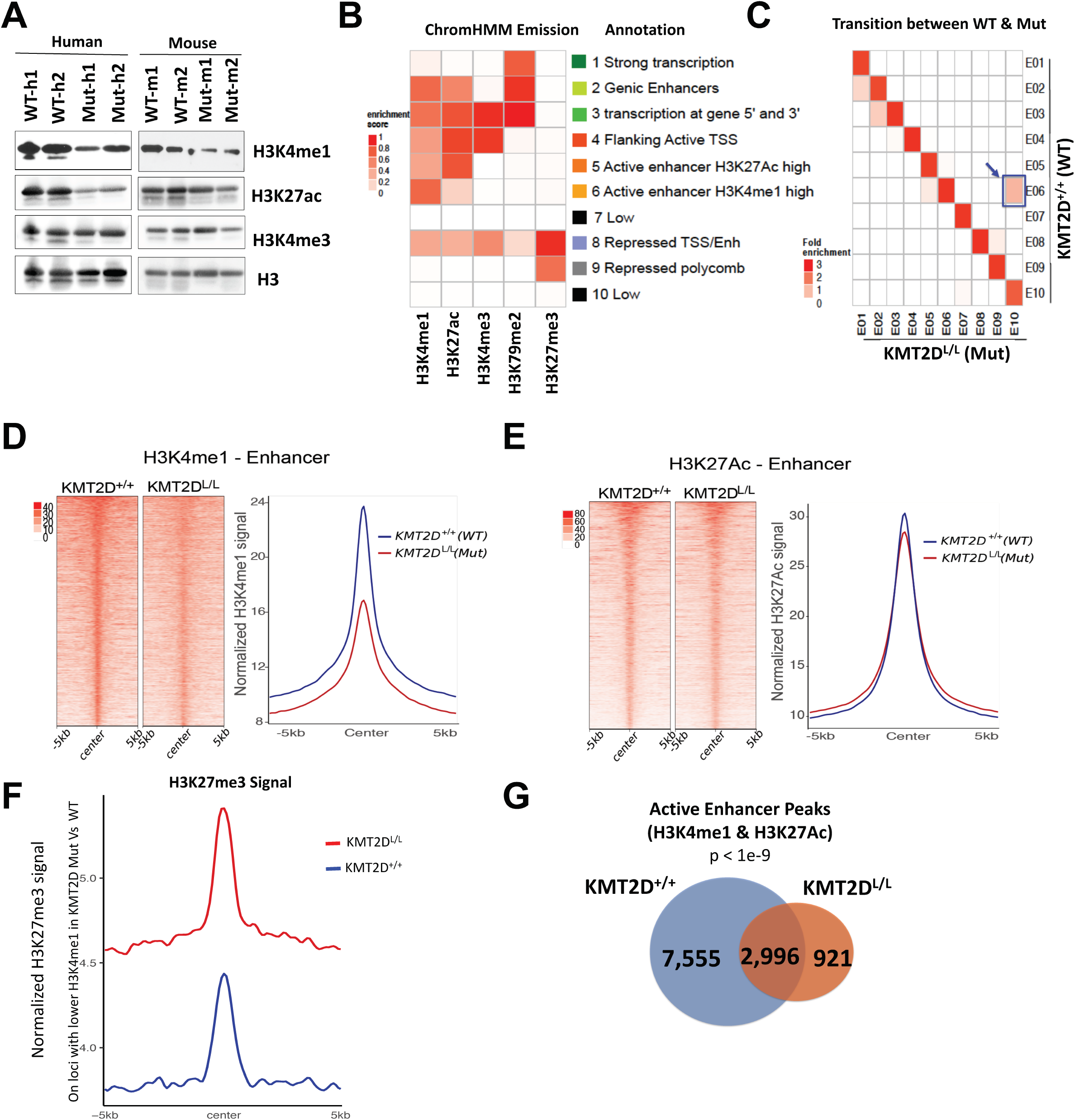
KMT2D loss is associated with loss of H3K4me1-marked enhancers. **A.** Western blot showing total H3K4me1, H3K27ac, H3K4me3, and H3 in KMT2D mutant (Mut-m1, Mut-m2, Mut-h1, Mut-h2) and WT (WT-m1, WT-m2, WT-h1, WT-h2) murine (right) and human (left) melanoma cells. **B.** Emission probabilities of the 10-state ChromHMM model on the basis of ChIP-Seq profiles of five histone marks (shown in X-axis). Each row represents one chromatin state and each column corresponds to one chromatin mark. The intensity of the color in each cell reflects the frequency of occurrence of that mark in the corresponding chromatin state on the scale from 0 (white) to 1 (red). States were manually grouped and given candidate annotations. **C.** Heat map showing the fold enrichment of chromatin state transitions between KMT2D mutant (KMT2D^L/L^) and WT (KMT2D^+/+^) samples for the 10-state model defined by the ChromHMM. Color intensities represent the relative fold enrichment. Blue box and arrow point to active enhancer state switch. **D-F.** Heat maps (left panels) and average intensity curves (right panels) of ChIP-Seq reads (RPKM) for H3K4me1 (**D**), H3K27ac (**E**) and H3k427me3 (**F**) at typical enhancer regions. Enhancers are shown in a 10kb window centered on the middle of the enhancer) in *iBIP;KMT2D^+/+^* and *iBIP;KMT2D^L/L^* melanoma tumors. Panel F shows signal for H3K27me3 on regions that lose H3K4me1 mark in KMT2D mutants in comparison to wild type tumors. **H.** Venn diagram showing unique or shared H3K4me1 and H3K27Ac co-enriched active enhancers sites in *iBIP;KMT2D*^+/+^ and *iBIP;KMT2D^L/L^* melanoma tumors

### Upregulated Insulin Growth Factor (IGF) Signaling Regulates Glycolysis in KMT2D Mutants

In order to understand how enhancer loss may lead to observed metabolic reprogramming, we overlapped changes in gene expression between KMT2D WT and mutant murine tumor-derived lines with the changes in active enhancer patterns. Of the 6980 active enhancer loci that display loss of intensity in KMT2D mutant tumors compared to WT, 1165 were located nearby (+/- 5Kb) to genes with decreased expression (**Figure 6A**). We found a significant association between loss of expression and loss of H3K4me1 patterns in nearby loci (**Figure S6A**). These genes were enriched for those involved in various phosphorylation mediated cell signaling events and are *bona fide* or putative tumor suppressors (**Figure S6B-C**). Of these, we focused on the IGF (Insulin Growth Factor) signaling pathway which is known to play major roles in regulating metabolic pathways (such as glycolysis) via activation of AKT(62) (**Figure S6B-C**). Indeed, we observed higher levels of pAKT (S473) and pIGF1R (Y1198) in KMT2D mutant murine and human lines (**Figure 6B**) as well as KMT2D mutant iBIP tumors (**Figure 6C**) suggesting aberrant activation of IGF-AKT-Glycolysis pathway. Examination of Reverse Protein Phase Array (RPPA) data from Cancer Cell Line Encyclopedia (CCLE) database(63) across all cancer types showed that KMT2D mutant cell lines (harboring functional driver mutations) showed higher levels of pS473 and pT308 forms of AKT compared to KMT2D wild type (and high expressing) lines (**Figure 6D**). Functional significance of activation of IGF1R signaling was further tested by treatment of cells with IGF-1R inhibitor (Linsitinib), which reduced expression of glycolysis genes in KMT2D mutant murine and human cell lines (**Figure 6E-F**). Importantly, treatment of KMT2D mutant murine and human cell lines with Linsitinib preferentially reduced the proliferation of KMT2D mutant cell lines both *in vitro* (**Figure 6G**) and *in vivo* (**Figure 6H-I**). This was recapitulated in the analysis of all cancer cell lines (from Sanger Cell Line database) for which linsitinib sensitivity data was available. Cells harboring KMT2D ‘functional’ driver mutations displayed significantly lower IC_50_ values for Linsitinib treatment compared to cells that harbor high levels of KMT2D (and have WT protein) (**Figure 6J**). Finally, Linsitinib treated tumors showed a drastic reduction in expression of glycolysis genes, proliferation marker Ki-67 and pAKT levels (**Figure S6C**). These data establish activation of IGF1R-AKT-Glycolysis axis in KMT2D-deficient cancer cells.

**Figure 6:**
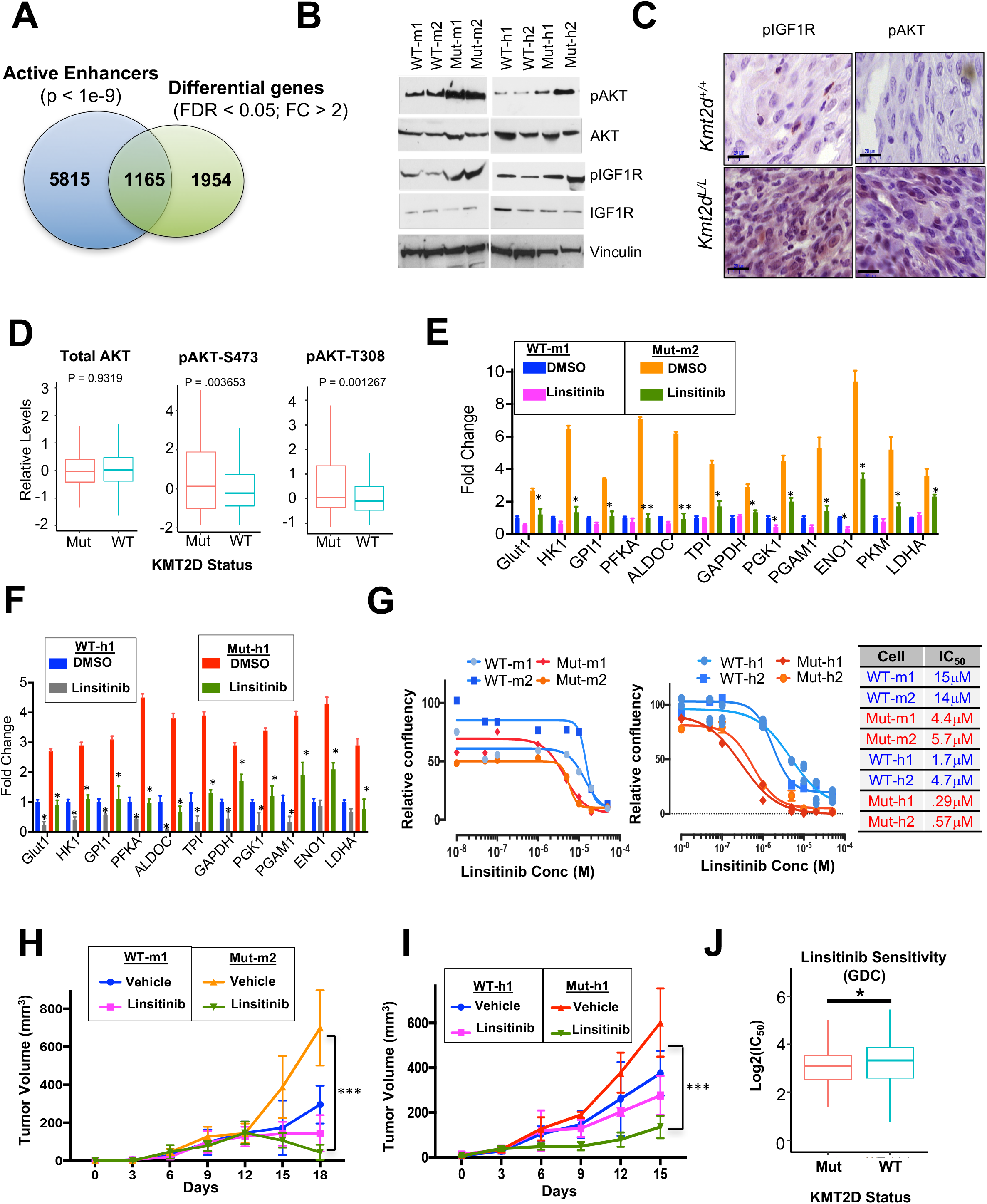
Aberrant activation of IGF and AKT signaling in KMT2D mutants confers sensitivity to IGFR inhibitor. **A.** Venn diagram showing the overlap between differentially expressed genes and lost active enhancer loci in KMT2D mutant (Mut-m2) melanoma cells compared to KMT2D WT (WT-m1). **B.** Western blot showing expression of pAKT, AKT, pIGF1R and IGF1R in KMT2D mutant and wild type murine (left panel) and human (right panel) cells. Vinculin is used as a loading control. **C.** Immunohistochemistry images for pAKT and pIGF1R demonstrating their overexpression in *iBIP;KMT2D*^+/+^ and *iBIP;KMT2D^L/L^* melanoma tumors. **D.** Box plot showing RPPA-based protein levels of AKT, pAKT (S473) and pAKT (T308) in Cancer Cell Line Encyclopedia (CCLE) cell lines (all cancer types) that harbor KMT2D functional mutations (n = 15) versus those that harbor wild type and high levels of KMT2D (n = 15; RPKM >= 10); t-test *p < 0.05, **p < 0.01. **E-F.** Bar graph showing relative expression pattern of 12 glycolysis enzyme genes (compared to 28S) in KMT2D mutant and WT murine (**D**) and human (**E**) cells treated with vehicle or Linsitinib (1μM) for 24 hours. T-test *p < 0.05, **p < 0.01. **G.** Growth curves for KMT2D mutant and WT murine (**F**) and human (**G**) melanoma cells treated with different concentrations of Linsitinib. Relative confluence at 96-hours post treatment are plotted and IC50 values are shown in the accompanying table. **H-I.** Line plot showing average tumor volumes for mice (n = 5 per group) injected with KMT2D mutant and WT murine (**H**) and human (**I**) melanoma cells and treated with Linsitinib (25mg/kg) or vehicle (30% PEG-400) every other day. **J.** Box plot showing Log(IC_50_) values for Linsitinib in the KMT2D mutant and KMT2D WT-high (n = 15; KMT2D RPKM ≥ 10) cell lines (all cancer types) from the GDC data (Sanger Cell Line Project).

### Loss of a Distal Enhancer of IGFBP5 in KMT2D Mutants Partially Regulates IGF Signaling and Expression of Glycolysis Enzyme Genes

We next searched for putative regulators of the IGF signaling that lose active enhancers and gene expression in KMT2D mutants specifically in order to identify those that may be responsible for the metabolic reprogramming phenotypes observed in KMT2D-deficient tumors. We focused on IGFBP5 as it is a known negative regulator of IGF1R signaling and acts as a tumor suppressor in melanoma by regulation of AKT and IGF1R signaling (62). We found the loss of H3K4me1 signals on proximal and distal enhancers associated with IGFBP5 in KMT2D mutant tissues (**Figures 7A**) while other IGFBPs did not show significant change (**Figure S7A**). Examination of Hi-C based higher order chromatin interaction data showed that IGFBP5 may be located in a TAD (Tandem Adjacent Domain) domain thus promoting interaction between this distal enhancer and IGFBP5 gene (**Figure 7B**). Consistently, IGFBP5 expression was also lost in KMT2D mutant murine and human cell lines (**Figure 7C**) while several other IGFBPs showed inconsistent patterns (**Figure S7B**). Consistently, IGFBP5 expression was significantly reduced in KMT2D mutant melanoma tumors (**Figure 7D and S7C**).

**Figure 7:**
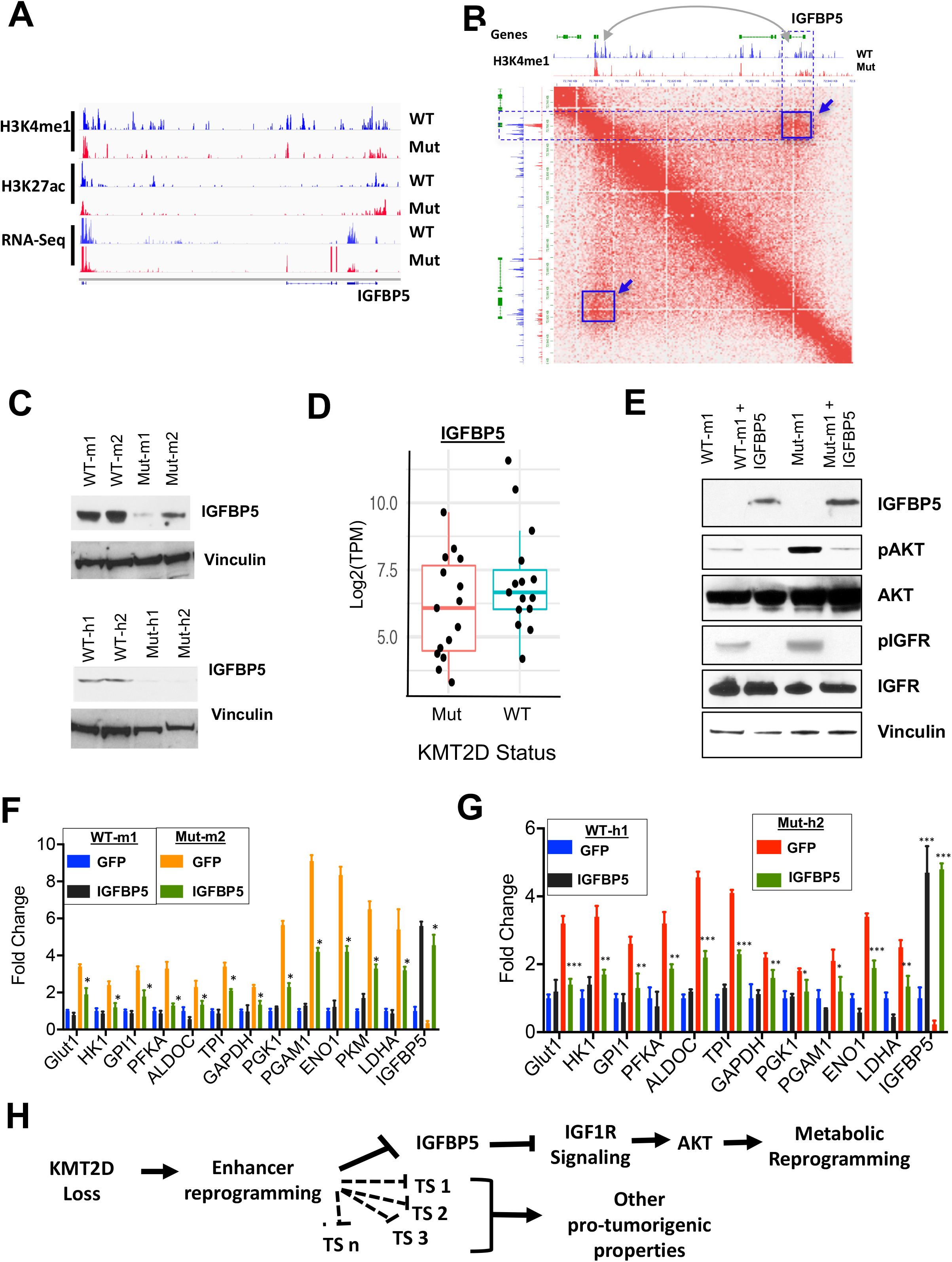
Loss of distal enhancers of IGFBP5 in KMT2D mutant cells partially contribute to its phenotypes. **A.** IGV snapshot showing RNA-seq, H3K27Ac, H3K4me1 ChIP-seq signal tracks for genomic locus harboring IGFBP5. Note the loss of blue peaks in region surrounding IGFBP5 gene. **B.** Interaction map from mouse ES cells showing IGFBP5-containing locus, demonstrating interaction between the IGFBP5 gene with proximal and distal enhancers. The highlighted blue off-diagonal interaction (box with arrow) points to a downstream enhancer that exhibits selective loss of H3K4me1 signal in KMT2D mutant samples. **C.** Western blot showing expression of IGFBP5 and Vinculin in KMT2D mutant and wild type murine (top panel) and human (bottom panel) cells. **D.** Box plot showing expression of IGFBP5 in the melanoma TCGA samples that harbor functional mutations (nonsense, frameshift or post4700aa) (n = 15) or WT copies for KMT2D (n = 15). **E.** Western blot showing expression of IGFBP5, pAKT, AKT, pIGF1R and IGF1R in KMT2D mutant and wild type murine control or IGFBP5 overexpressing cells. Vinculin is used as a loading control. **F-G.** Bar graph showing relative expression pattern of 12 glycolysis enzyme genes (compared to 28s) in KMT2D mutant and WT murine (**F**) and human (**G**) cells overexpressing empty vector or IGFBP5. Standard et-test *p < 0.05, **p < 0.01. **H.** Model of KMT2D function in melanoma. Our data suggest that KMT2D loss leads to enhancer reprogramming on tumor suppressor genes including IGFBP5 which control various pathways such as IGF1R signaling that leads to activation of AKT and rewires metabolic pathways.

Functional significance of loss of IGFBP5 was further tested by overexpression of IGFBP5 in KMT2D WT and mutant murine cells. IGFBP5 overexpressing murine melanoma cells showed lower levels of IGF1R and AKT phosphorylation (**Figure 7E**). Consistently, reduced expression of glycolysis genes were observed preferentially in in KMT2D mutant murine and human cells compared to their wild type counterparts (**Figure 7F-G**). Taken together, the data presented in this manuscript establish a model of KMT2D function in cancer where KMT2D acts as a tumor-suppressor by enhancer reprogramming on tumor suppressor genes such as IGFBP5 that regulate key pathways such as IGF1R signaling leading to metabolic rewiring (**Figure 7H**).

## DISCUSSION

Although the somatic loss of function mutations in KMT2D are observed across a large number of malignancies (37–52), it is unclear why these mutations are selected over course of tumor evolution. Our study suggests that enhancer reprogramming via KMT2D loss may rewire metabolic pathways for increased energy and biomass needs of cancer cells. We observed drastic deregulation of multiple metabolic pathways in KMT2D mutant melanomas in both human and murine systems. Consistently, we observed preferential dependence of KMT2D mutant cells’ growth on glycolysis in comparison to wild type cells. Glycolysis pathway serves as a central node for various needs of a proliferating cells (64). It is required for a small fraction of energy needs (2 ATPs per cycle), and, more importantly, for production of biomass needed for cell doubling. For example, glucose-6-phosphate provides gateway to nucleotide biosynthesis and dihydroxyacetone phosphate acts as a starting substrate for lipid biosynthesis pathway. Increased pyruvate production due to high glycolysis provides substrate for the OxPhos pathway (to generate 36 ATPs), which is also upregulated in the KMT2D mutant cells, thereby leading to enhanced ATP production. Finally, 3-phosphoglycerate and other OxPhos metabolites provide substrate for amino acid biosynthesis. Therefore, upregulated glycolysis in KMT2D mutant cells is critical for many different needs to enhance tumorigenesis.

Although we show an important role for glycolysis, many other metabolic pathways such as oxidative phosphorylation and fatty acid metabolism are also highly upregulated in KMT2D mutant cancers. The publically available CRISPR screening platform, Achilles (65), suggests a dependency of KMT2D mutant melanomas on specific genes in these other metabolic pathways which need further exploration. Indeed, a recent study suggested enhanced fatty acid metabolism in pancreatic cancer (53). Interestingly, the dependence of SMARCA4 mutant cancers on the blockade of oxidative phosphorylation was recently demonstrated (66) in lung cancer. Together, these studies suggest the need of in-depth clinical investigation on potential usage of metabolic inhibitors in patients with mutations in SWI/SNF and MLL complexes.

Our data along with the accompanying manuscript by Alam et al (See supplementary material for the manuscript) show the dependence of KMT2D mutant cancers on glycolysis and, critically, will inform future clinical studies testing potent glycolysis-blocking inhibitors in this genetic context. Our data also suggest the potential use of IGF receptor blocking molecules such as Linsitinib, which is being tested in clinical trials (67), in the KMT2D mutant patient population. However, further work may be needed to better stratify the functional driver mutations in KMT2D because it is a long gene and some of the observed somatic mutations may be passenger events, especially in cancers with high mutation burden such as melanoma and lung cancers (68). In addition to mutations, KMT2D expression levels may also need consideration while stratifying patients for such therapies as large number of metastatic and primary melanomas show little to no expression of KMT2D.

KMT2D is a member of the COMPASS complex which is thought to be critical for depositing H3K4me3(69,70). Furthermore, some studies, such as one by Dhar et al (4) suggest a role of KMT2D in H3K4me3 regulation. However, several other studies suggest KMT2D to be a major regulator of the H3K4me1 mark which marks poised enhancers (17,19,20,58,69,71,72). In a subset of enhancers H3K4me1 recruits CBP/p300 enzymes in turn activation their target genes (18), however, complete understanding this mode of active enhancer regulation is still lacking. Our data suggest that KMT2D is a major regulator of H3K4me1 in melanoma. Intriguingly, since we did not see drastic changes in H3K27ac it is possible that other histone acetylations (than H3K27ac) may be involved in enhancer activation in KMT2D mutant melanomas. Indeed, evidence for roles the of other histone acetylations in enhancer activation has been previously demonstrated (73). Our previous study also showed drastic losses of chromatin states harboring multiple different histone acetylation and H3K4me1/2/3 in early stages of tumorigenic transition in melanoma (33).

KMT2D was previously shown to promote proliferation in NRAS-mutant patient-derived xenograft melanoma cells(74), which contrasts with our findings. There are several possible explanations for the discrepancy between the results of these two studies. First, our study examines the role of KMT2D in BRAF mutant conditions whereas other Bossi et al(74) focused on NRAS mutant melanomas. It is possible that KMT2D has distinct functions in these two genetic contexts, perhaps influenced by contexts-specific and differing interaction partners or post-translational modifications. Secondly, given the dependency on metabolic pathways as well as upregulation of immune mechanisms observed in melanoma, the discrepancy in the two studies could reflect the use of model systems. Our study provides evidence for tumor-suppressor roles using an immune-competent genetically engineered mouse model whereas Bossi et al used immunodeficient mice. In addition, the growth medium for PDXs or cell lines is critical because of the high glycolytic rates of KMT2D-deficient cells. Available glucose in the media is utilized rapidly likely leading to suppression of cell cycle. Indeed, the establishment of KMT2D mutant cells from GEMM model (*iBIP:KMT2D^L/L^*) tumors required a repeated change of DMEM media with high glucose every 3-4 hours (**Figure S3E-F**). Future studies with GEMM models of KMT2D in NRAS mutant conditions will be needed to address some of these possibilities.

Overall, our study provides evidence for the dependency of the KMT2D mutant melanomas on glycolysis and the IGF pathway via enhancer reprogramming. These results suggest a potential and novel therapeutic strategy in the patients with melanoma harboring mutations in this epigenetic regulator.

## METHODS

All methods are available as supplementary information.

## Supporting information

Genes and shRNAs included in the screen

Pathways enriched in KMT2D mutant tumors

List of reagents

## AUTHOR CONTRIBUTIONS

M.M. and E. Z. K. planned and carried out experiments, analyzed data, prepared figures, and wrote the manuscript. M.T. performed the bioinformatics analysis of RNA-seq, ChIP-seq, TCGA, and CCLE/GDC data. H.A. contributed to experimental design, provided reagents and helped with preparation of figures. L.Y. and H.Y. performed metabolic experiments and helped with data analysis. G.H. and. L.W. contributed to the TCGA data analysis. A.T.R., C.T. and. S.B.A. helped with informatics analysis. S. Sarkar and E.O. contributed to study design and provided help with experimentation. S. Sharma, M.W., N.S.S. M.D. N.Z., T.S., A.S., J.B.A. N.E.A. provided technical help. B.G. contributed to development of mouse mode. S. J. provided support for mouse colony maintenance and experimental help. E.Q.C. contributed to IHC experiments and analysis. A.L. provided guidance on mouse tumor pathology. Y-H.L. and F.M. provided reagents. K.R. conceived and designed the study, performed experiments, evaluated data, made figures and wrote the manuscript.

## ACKNOWLEDGEMENTS

We thank Marcus Coyle, Curtis Gumbs, SMF core at MDACC for sequencing support. We thank Yanping Cao, Jill Garvey, Ibarra Ivonne and Kun Zhao for mouse colony maintenance. The work described in this article was supported by grants from the National Institutes of Health (CA160578 to K. R.; CA157919, CA207109, and CA207098 to M.G.L. and CA016672 to SMF Core), American Cancer Society (RSG-15-145-01-CDD to F.M.), Center for Cancer Epigenetics at MDACC (K.R.) and MD Anderson Cancer Center (Start-up funds to K.R.). The following fellowship support is acknowledged: K. R. (Charles A. King postdoctoral fellowship) and H.A. (Odyssey Fellowship at MD Anderson Cancer Center). The histology work were performed at the Histopathology Core Lab at the UT MD Anderson Cancer Center supported by the NIH National Cancer Institute (P30CA016672).

