## Supplementary Methods, Figures and Legends for "Enhancer Reprogramming Confers Dependence on Glycolysis and IGF signaling in KMT2D Mutant Melanoma"

#### SUPPLEMENTARY INFORMATION

#### METHODS

##### Cell culture, stable cell generation and inhibitor treatment

HMEL-BRAF$^{V600E}$, A375, RPMI-7951, WM115 and WM266-4 cells were maintained in standard tissue-culture conditions in DMEM media (high glucose) with 10% FBS. SKMEL-24 (Mut-h1) and WM278 (Mut-h2) cells were maintained in the recommended media, except all assays were performed in the same media as for A735 (WT-h1) and RPMI-7951 (WT-h2). Mouse tumor cell lines 300 (WT-m1), 5770 (WT-m2), 3417 (mut-m1) and 3418 (Mut-m2) were isolated from melanoma tumors by digestion in RPMI media (Sigma) using collagenase (2mg/ml) (Gibco) and dispase (4mg/ml) (Gibco). Single cell suspension was generated using MACS homogenizer (Milteny Biotec) following manufacturer's mTumor protocol. Cells were plated in DMEM with high glucose (Sigma/Gibco) and Glutamax (Sigma/Gibco) and replenished every 4 hours. Once cultures were stable, cells were maintained in DMEM with high glucose and Glutamax. Stable lines expressing shRNAs and ORFs were established by standard lentiviral mediated transduction. All cells were routinely tested for mycoplasma by mycoAlert kit (Lonza) or by PCR.

##### Inhibitor treatment experiments

For inhibitor treatment experiments, cells were seeded in 96-well plates at a density of 500 cells per well and treated 24-hours later with specific inhibitors. Linsitinib (SelleckChem), 2-Deoxy-D-Glucose (2-DG, Sigma) and Lonidamine (Sigma) were dissolved in DMSO and diluted in cell culture media for treatment. Cell density was measured by imaging every 3-4 hours using an IncuCyte live cell analysis system (Essen Bioscience). Once treated cells reached confluence (90-100%), they were stained with crystal violet and destained in 10% acetic acid. Absorbance was read at 590nM and background-corrected readings for each condition were normalized to vehicle treatments which were plotted to obtain the growth curves. $IC_{50}$ values were calculated using GraphPad Prism.

**RNAi screen**

For the screen, 475 shRNAs against 95 chromatin modifiers (**Supplemental Table S1**) in pLKO.1 vector were obtained from the Broad Institute/Harvard Medical School. Lentiviruses were produced in HEK-293T cells in a 6-well format using pDelta8.2 and pMD2.G (addgene). Viruses for 25 shRNAs (5 shRNAs for each of 5 genes selected at random) were pooled together and the mixture was used to transfect 1 million cells, which were subsequently grafted intradermally into the flanks of NCR-NUDE mice. All mice experiments were performed per Institutional Animal Care and Use Committee (IACUC) guidelines. Mice were monitored weekly for tumor growth and sacrificed when tumor size reached 2 cm in one dimension. At sacrifice, tumors were harvested and genomic DNA prepared. The segment of pLKO.1 containing shRNA sequence was amplified by PCR and then sequenced to identify the integrated shRNA. Genes corresponding to these shRNAs were noted as hits from the primary screen. For validation experiments, stable HMEL-$BRAF^{V600E}$ cells were generated using individual shRNAs, with multiple shRNAs targeting each gene from primary hits. Cell lines generated from the two best shRNAs demonstrating the best knockdown of the corresponding gene (and with at least 50% knockdown) were then injected intradermally in the flanks of NCR-NUDE mice (1 million cells/injection). Mice were monitored for tumor formation and growth.

**Mouse experiments**

All mice were kept in specific pathogen free vivarium at the MD Anderson Cancer Center mouse facilities. Mice were fed commercial rodent diet (PicoLab Rodent diet 5053 from Labdiet) and water *ad libitum*. All animal studies and procedures have complied with ethical regulations and all protocols pertaining to animal studies and procedures have been approved by The University of Texas MD Anderson Cancer Center Institutional Animal Care and Use Committee (IACUC).

***Xenograft experiments***

All mice experiments were performed per Institutional Animal Care and Use Committee (IACUC) guidelines. Mice were injected with 5 million cells in one flank each and monitored every other day for tumor growth. When tumor size reached 0.5 cm in one mouse in each treatment arm, all mice were injected with 2-DG (500mg/kg dissolved in PBS) or Linsitinib (25mg/kg dissolved in 30% PEG-400) via intraperitoneal route. Tumor volume was measured every other days.

***Genetically engineered mouse model***

KMT2D mutant mice were obtained from Dr. MinGyu Lee(1). These mice are engineered with Lox sites flanking exons 16 and 20, resulting in loss of KMT2D protein as previously described(1). KMT2D mutant mice were crossed with iBIP mice(2) (iBIP = Tyr-Cre$^{ERT2}$, Rosa26-rtta, TetO-BRAF$^{V600E}$, PTEN$^{L/L}$, INK/ARF$^{L/L}$; mixed genetic background of FVB and B6) to generate *iBIP;KMT2D*$^{L/+}$ genotype containing mice (as assessed by genotyping for all alleles). The *iBIP;KMT2D*$^{L/+}$ male and female mice were mated to generate *iBIP;KMT2D*$^{+/+}$, *iBIP;KMT2D*$^{L/+}$, and *iBIP;KMT2D*$^{L/L}$ genotype containing cohorts. These mice were treated with doxycycline systemically (2mg/ml in 40mg/ml of water) by feeding (*ad libitum)* and 4-OHT (1$\mu$M) was applied on the ears to generate auricular tumors. Mice were observed twice a week for tumor formation and upon tumor appearance, tumor growth was measured every other day. Tumors were harvested by excision of the lesion and digested for generation of WT-m1, WT-m2, Mut-m1 and Mut-m2 cell lines.

**RNA-Seq analysis of murine tumor cells**

Strand specific libraries were constructed using the ScriptSeq Kit (Epicenter/Illumina). RNAseq data were processed by pyflow-RNAseq(3), a snakemake based RNAseq pipeline. Raw reads were mapped by STAR(4), RPKM normalized bigwigs were generated by *deeptools*(5), and gene counts were obtained by *featureCount*(6). Differential expression analysis was carried out using *DESeq2*(7). Gene set enrichment analysis(GSEA) was done using the GSEA tool(8) from Broad Institute. The pre-rank mode was used. The signed fold change *–log10(pvalue) metric was used to pre-rank the genes.

**ChIP-Seq**

Chromatin immunoprecipitation was performed as described earlier(9) with optimized shearing conditions and minor modifications for melanocytes. The antibodies used were: H3K4me1 (Abcam ab8895), H3K27ac (Abcam ab4729), H3K4me3 (Abcam ab8580), H3K79me2 (Abcam ab3594), H3K27me3 (Abcam ab6002). ChIP-seq data were quality controlled and processed by pyflow-ChIPseq(10), a snakemake(11) based ChIPseq pipeline. Briefly, raw reads were mapped by bowtie1(12) to hg19. Duplicated reads were removed and only uniquely mapped reads were retained. RPKM normalized bigwigs were generated by deep tools(5) and tracks were visualized with IGV(13). Peaks were called using macs1.4(14) with a p-value of 1e-9. Chromatin state was called using ChromHMM(15) and the emission profile was plotted by ComplexHeatmap(16). Heatmaps were generated using R package *EnrichedHeatmap*. ChIP-seq peaks were annotated with the nearest genes using ChIPseeker (17). Super-enhancers were identified using ROSE(18) based on H3K27ac ChIP-seq data.

**Chromatin state analysis**

ChromHMM(15) was used with default parameters to derive genome-wide chromatin state maps for all cell types. Input data was binarized using ChromHMM's BinarizeBed method (15) with a p-value cutoff of $1e^{-4}$. Chromatin state models were learnt jointly on all data for all 5 histone marks

(H3K4me1, H3K4me3, H3K27ac, H3K79me2 and H3K27me3) from WT-m1 and Mut-m2 tumor cells and a model with 10 states was chosen for detailed analysis. Chromatin state segmentations of WT-m1 and Mut-m1 were produced subsequently by applying this model to the original binarized, quantile normalized or downsampled chromatin data from these cell types.

**TCGA RNA-Seq data analysis**

TCGA melanoma (SKCM) RNAseq raw counts were downloaded using TCGAbiolinks(19). The mutation MAF files were downloaded with TCGAbiolinks as well. Mutation status of KMT2D was inferred from the MAF files. Ten SKCM primary tumor samples with wild type copies of KMT2D and expressing high levels and 10 SKCM primary tumor samples with mutant KMT2D (see supplementary data for samples included in the analysis) were compared using DESeq2, the signed fold change *–log10(pvalue) metric was used to pre-rank the gene list and for GSEA pre-rank analysis.

For the pan cancer analysis, TCGA tumor samples were grouped based on KMT2D gene expression and mutation status: KMT2D mutation free group are samples with high KMT2D expression (among the top quantile) and no somatic mutation; KMT2D mutated group are samples with low KMT2D expression (falling into the bottom quantile) and have either nonsense mutations or missense mutations with the amino acid 4,700. Six tumor types (BLCA, urothelial bladder carcinoma; CESC, squamous cell carcinoma and endocervical adenocarcinoma; HNSC, head and neck squamous cell carcinoma; LUSC, lung squamous cell carcinoma; UCEC, uterine corpus endometrial carcinoma; STAD, stomach adenocarcinoma) had adequate sample size (≥10) to be included for differential gene expression analysis and pathway enrichment analysis. TCGA normalized RNAseq read count were processed by Wilcoxon test to identify differentially expressed genes (DEGs) across KMT2D mutation status groups. A cut-off of gene expression fold change of ≥ 2 or ≤ -0.5 and a FDR q-value of <0.05 were applied to select the most differentially expressed genes. A ranked list of genes was generated based on the product of

Wilcoxon test FDR q-values and log2 fold change for all coding genes and processed by Gene Set Enrichment Analysis (GSEA)(8) against the curated gene sets from Molecular Signature Database (MSigDB)(20) to identify significantly enriched signaling pathways.

**Immunohistochemistry**

Tumors were fixed in formalin for 24 hours, paraffin embedded, sectioned and stained according to standard procedures. Briefly, endogenous peroxidases were inactivated by 3% hydrogen 673 peroxide. Non-specific signals were blocked using 3% BSA, 10% goat serum in 0.1% Triton X-100. After antigen retrieval in citrate buffer, slides were stained using respective antibodies overnight at $4^{o}$C, [See **Supplementary Table S3** - ENO1 (Proteintech, #11204-1-AP), PGK1 (Proteintech #17811-1-AP), PGAM1 (Proteintech, #16126-1-AP), Tyrosinase (Abcam, ab738), pAKT (Cell Signaling, #9271), pIGF1R (Abcam, ab39398)]. After overnight incubation, the slides were washed and incubated with secondary antibody (HRP-polymers, Biocare Medical) for 30 min at room temperature. The slides were washed three times and stained with DAB substrate (ThermoFisher Scientific). The slides were then counterstained with haematoxylin and mounted with mounting medium.

**Inducible ectopic expression of KMT2D**

Full length KMT2D was cloned from KMT2D overexpression vector (gift from Dr. Laura Pasqualucci at Columbia University) into the pInducer20 doxycycline inducible lentiviral vector (Addgene 44012). Lentivirus was produced using standard virus production methods by co-transfecting target and packaging plasmids (psPAX2 – Addgene12260 and pMD2.G- Addgene 12259) into HEK293T cells. Cell lines were then transduced with 0.45uM filtered and ultracentrifuge concentrated viral particles with Polybrene (8µg/ml). After 16 hours of transduction, media was changed into fresh regular growth media, and 48 hours later selection started using G418 (0.2-0.6µg/ml). After selection was complete in 120 hours, cells were termed stably transduced. KMT2D expression was induced for 72 hours with doxycycline 2µg/ml.

**Whole Cell Extracts, Acid Extraction and Western Blotting**

Cells were harvested in media, washed with PBS, and pelleted. Cell pellets were dissolved in RIPA buffer (25mM Tris PH8, 150mM NaCl, 01% SDS, 0.5% Sodium Deoxycholate, 1% Triton X-100, Protease and phosphatase inhibitor cocktail; bought from Boston bioproducts) and incubated for 30' on ice before brief sonication followed by centrifugation to remove debris. Supernatant was collected, protein measured using Bradford assay and equal amounts were loaded on the 4-12% SDS-PAGE gel (Invitrogen). Proteins were transferred to a Nitrocellulose or PVDF membrane which was then blocked in Odyssey Blocking buffer (LiCOR) and incubated with primary antibody overnight in the same buffer. Blots were then washed and probed with HRP-labelled secondary antibodies (Pierce) and developed using a X-ray film (Phenix). For histone marks, we incubated cell pellets in Triton Extraction buffer (PBS containing 0.5% Triton X 100 (v/v), 2 mM phenylmethylsulfonyl fluoride (PMSF), 0.02% (w/v) $NaN_3$) for nuclei isolation. Nuclei were subjected to histone extraction by overnight incubation in 0.2N HCl (with protease and phosphatase inhibitors) followed by centrifugation. Rest of the western blotting was done as explained above. Antibodies used were (also listed in **Supplementary Table S3**): IGFBP5 (Proteintech, #55205-1-AP), pAKT (Cell Signaling, 9271) Total AKT (CST, #4691), Total IGF-1R (Cell signaling, #9750). Histone antibodies are same as used for ChIP experiments.

**Metabolomics via Selected Reaction Monitoring Tandem Mass Spectrometry**

One 15 $cm^2$ plate of cells (~10–15 million) per sample was extracted with 80% methanol (−80°C) for 15 min. Dried metabolite pellets were resuspended in 20 µL of LC/MS-grade water, and 5 µL aliquots were injected for targeted LC/MS/MS on a 5500 QTRAP hybrid triple-quadrupole mass spectrometer coupled to a Prominence ultrafast liquid chromatography (UFLC) system from 287 selected reaction monitoring (SRM) transitions with positive/negative polarity switching. Samples were separated on a 4.6 mm i.d. × 100 mm Amide XBridge hydrophilic interaction liquid chromatography (HILIC) column at 360 µL/min starting from 85% buffer B (100% ACN) and moving to 0% B over 16 min. Buffer A was 20 mM $NH_4OH$/20 mM $CH_3COONH_4$ (pH=9.0) in 95:5 water/ACN. Q3 peak areas were integrated by use of MultiQuant 2.1 software (AB/SCIEX). MetaboAnalyst 2.0 (http://www.metaboanalyst.ca) was used to normalize data. All metabolite samples were prepared as biological triplicates.

**RT-qPCR**

RNA was isolated using RNeasy kit (qiagen) or Trizol (Thermo Fisher) reagent using manufacturer's protocol. cDNA was prepared using SuperScript III first strand synthesis kit (Thermo Fisher) using 2micrograms of RNA and manufacturer's protocol. Quantitative PCR was performed using QuantiTect Sybr Green PCR kit in Stratagene's Mx3000p system. Primers used are listed in **Supplementary Table S3**.

**Tissue Microarray**

The tissue microarray containing 100 samples (62 cases of primary melanoma, 20 cases of metastatic melanoma, and 18 nevi) was obtained from US Biomax. The staining for KMT2D antibody (Sigma, Prestige) was performed at the immunohistochemistry core at MD Anderson Cancer Center. Two pathologists read the TMA and consensus scores were assigned to each sample.

**DATA AVAILABILITY STATEMENT**

All RNA-Seq and ChIP-Seq data are available at GEO accession number GSE116921.

#### CODE AVAILABILITY

All codes used to generate the ChIP-Seq data are available at https://github.com/crazyhottommy/pyflow-ChIPseq. Rest of the codes are available upon request.

#### SUPPLEMENTARY FIGURE LEGENDS

**Supplementary Figure S1: Validation of epigenetic regulators identified through RNAi screen identified 8 epigenetic regulators as potential tumor suppressors in melanoma.**

**A.** Kaplan-Meier curve showing tumor-free survival of mouse cohorts orthotopically injected with 1 million HMEL-BRAF$^{v600E}$ cells transfected with pooled shRNAs from primary screen. This figure shows data from pools that did not significantly accelerate tumorigenesis. Three negative control pools (shLuc, shGFP and shNT) are shown. Cells from each pool were injected in 10 mice each and tumor formation was monitored over 25 weeks.

**B-I.** Bar graph showing relative levels of indicated genes in the HMEL-BRAF$^{V600E}$ and WM115 cells harboring lentivirally integrated shRNAs for that gene or GFP. Y-axis represents fold change of the gene expression compared to 28s and normalized to control shRNA samples. Standard t-test *p < 0.05, **p < 0.01, ***p < 0.001

**J-P.** Kaplan-Meier curve showing tumor-free survival of mouse cohorts orthotopically injected with WM115 cells stably expressing shRNAs against KDM1A (**J**), APOBEC2 (**K**), HDAC6 (**L**), KMT2F (**M**), SETD4 (**N**), KAT4 (**O**) and KDM5B (**P**). Mantel-cox test *p < 0.05, **p < 0.01, and ***p<0.001.

**Q.** Representative images for invaded cells from Boyden chamber assay of HMEL-BRAF$^{V600E}$ cells harboring lentivirally integrated shRNAs for GFP (control) or APOBEC2, HDAC6, KDM5B, KDM1A, SETD4, KMT2F or KAT4. #1 and #2 represent the duplicates for the experiment.

**Supplementary Figure S2: Validation of KMT2D as a tumor-suppressor in melanoma.**

**A.** Schematic of KAT4 protein showing missense mutations seen across all melanoma studies deposited in cBio portal. Green filled circles denote missense mutations whereas black filled circles represent truncating mutations Colored boxes within KAT4 schematic show different protein domains.

**B.** Dot plot showing relative KMT2D mRNA levels between nevi and primary melanomas (from

Reference 21) (left panel) and between primary and metastatic melanomas (from Reference 22) (right panel).

**C, D, F.** Graph showing tumor volume of HMEL-BRAF$^{V600E}$ cells (clonal variant 1 in panel **C** and clonal variant 2 in panels **D** and **F**) harboring either control (shNT) or KMT2D shRNAs (shKMT2D-1 in panel **d** and KMT2D-2 in panel **f**).

**E, G.** Kaplan-Meier curve showing tumor-free survival of mouse cohorts orthotopically injected with HMEL-BRAF$^{V600E}$ cells (clonal variant 2) harboring either control (shNT) or KMT2D shRNAs (shKMT2D-1 in panel **e** and KMT2D-2 in panel **g**).

**H.** Relative soft agar colon formation ability of HMEL-BRAF$^{V600E}$ or WM266-4 cells harboring control or 2 different KMT2D shRNAs (shKMT2D-1 and shKMT2D-2). Unpaired t-test *$p < 0.05$, **$p < 0.01$, and ***$p<0.001$.

**I.** Representative images and quantification of invaded HMEL-BRAF$^{V600E}$ cells stably transfected with control or KMT2D shRNAs in Boyden Chamber assay. Unpaired t-test *$p < 0.05$, **$p < 0.01$, and ***$p<0.001$.

**J.** Images of H&E stained and Tyrosinase stained (standard Immunohistochemistry) (40X) melanoma tumors from *iBIP;KMT2D*$^{+/+}$ and *iBIP;KMT2D*$^{L/L}$ mice. Scale bar is 20mM.

**K.** Images of **i**mmunofluorescence for KMT2D in WT-m1, WT-m2, Mut-m1, Mut-m2, Mut-m2 + dox (hKMT2D), WT-h1, WT-h2, Mut-h1, Mut-h2, and Mut-h1 + dox (hKMT2D) cells for KMT2D. Nuclear staining is shown by DAPI staining (shown in grayscale).

**Supplementary Figure S3: KMT2D mutant cells induce energy metabolism pathways and depend on glucose availability.**

**A-B.** Top 5 GO terms (**a**) and all significant HALLMARK terms for downregulated genes (FDR < 0.05, FC >2) (**b**) between KMT2D mutant murine cells and KMT2D wild type cells by total RNA-Seq analysis.

**C.** All significant HALLMARK pathway terms in differentially expressed genes between KMT2D mutant (carrying truncation, frameshift and post4700aa missense) and wild type human primary melanomas from the TCGA data.

**D.** Enrichment plots for HALLMARK glycolysis pathway in differentially expressed genes between between KMT2D mutant (carrying truncation, frameshift and post4700aa missense) and wild type human tumors from six different tumor types in TCGA data where n of mutant samples > 10.

**E-F.** Growth curves for WT-m1, WT-m2, Mut-m1 and Mut-m2 cells (**E**) and WT-h1, WT-h2, Mut-h1, Mut-h2 cells in DMEL high glucose (4g/L) or low glucose (1g/L) media. X-axis represent hours post plating and Y-axis shows percent confluence.

**Supplementary Figure S4: Inhibition of glycolysis preferentially impacts KMT2D mutant cells.**

**A-B.** Growth curves for KMT2D mutant and WT murine (**A**) and human (**B**) melanoma cells treated with varying concentrations of Lonidamine. Relative confluence at 96-hours post treatment are plotted and IC50 values are shown in the accompanying table.

**C-D.** Bar plot showing $IC_{50}$ values for Pomhex upon treatment of WT-m1, WT-m2, Mut-m1, Mut-m2 (**C**) or WT-h1, WT-h2, Mut-h1, Mut-h2 (**D**) cells in high glucose (4g/L) or low glucose (1g/L) DMEM media.

**E-H.** Line plot showing tumor volumes for individual mice (n = 5 per group) treated with 2-DG (500mg/kg) or vehicle every other day. Individual panels show data from mice injected with murine KMT2D WT (WT-m1) (**E**), murine KMT2D mut (Mut-m2) (**F**), human KMT2D WT (WT-h1) (**G**) and human KMT2D mutant (Mut-h1) (**H**) melanoma cells treated with vehicle or 2-DG.

**I.** Growth curves for KMT2D mutant and WT murine melanoma cells treated with varying concentrations of IACS-10759, an inhibitor of oxidative phosphorylation. Relative confluence at 96-hours post treatment are plotted and IC50 values are shown in the accompanying table.

**Supplementary Figure S5: Levels of H3K4me3, H3K27me3 and H3K79me2 in KMT2D WT**

**and mutant melanoma tumors.**

**A.** Western blot showing total H3K4me1 and H3 in KMT2D mutant (Mut-m2, Mut-h1) and rescue with hKMT2D (+dox) murine (left) and human (right) melanoma cells.

**B-C**. Heat map (left panels) and average intensity curves (right panels) for super-enhancer peaks based on H3K4me1 (**B**) and H3K27ac (**C**) ChIP-Seq data peak in *iBIP;KMT2D*$^{+/+}$ and *iBIP;KMT2D*$^{L/L}$ melanoma tumors.

**D-F.** Heat map (left panels) and average intensity curves (right panels) derived from the ChIP-Seq reads (RPKM) for H3K27me3 (**D**), H3K4me3 (**E**) and H3K79me2 (**F**) at enriched promoters in 10kb window centered on the middle of the peak in *iBIP;KMT2D*$^{+/+}$ and *iBIP;KMT2D*$^{L/L}$ melanoma tumors. Signals show enrichment in regions that enriched for these marks in respective type of tumor.

**G.** Pathway analysis for active enhancer (H3K4me1 and H3K27ac overlap) peaks specific to KMT2D mutant compared to KMT2D wild type tumors.

**Supplementary Figure S6: IGF signaling and aberrantly activates glycolysis in KMT2D mutant melanoma tumors.**

**A.** Heat maps showing differentially expressed genes and associated H3K4me1 signals in the vicinity as noted. R1, R2 and R3 refer to three replicates for WT-m1 and Mut-m2 cells.

**B.** All significant GO terms for genes which are differentially expressed and harbor loss of H3K4me1 mark at associated loci in KMT2D mutant melanoma cells compared to WT tumor cells.

**C.** Snapshots of IGV viewer for H3K4me1, H3K27ac and RNA-Seq signals on genomic loci surrounding various *bona fide* or putative tumor suppressor genes in KMT2D wild type or mutant tumors.

**D.** Images of H&E stained tumor or those stained for the product of three glycolysis genes (ENO1, PGK1 and PGAM1) or pAKT in xenograft tumors of Mut-m2 cells treated with vehicle or Linsitinib (20mg/kg). Magnification is shown below the images. Bar shows 20$\mu$M in all cases except H&E

images where it represents 50μM.

**Supplementary Figure S7: Loss of enhancer activity at IGFBP5 mediates activation of IGF signaling and aberrantly activates glycolysis in KMT2D mutant melanoma tumors.**

**A.** IGV snapshot showing RNA-seq, H3K27Ac, H3K4me1 ChIP-seq signal tracks for genomic locus associated with IGFBP proteins.

**B.** Bar graph showing relative expression levels of indicated IGFBP genes in murine KMT2D mutant (Mut-m1 and Mut-m2) and WT (WT-m1 and WT-m2) cells. Y-axis represents fold change of the gene expression compared to 28S and normalized to control shRNA samples. Standard t-test $*p < 0.05$, $**p < 0.01$, $***p < 0.001$

**C.** Box plot showing expression of IGFBP genes in the melanoma TCGA samples that harbor functional mutations (nonsense, frameshift or post4700aa) (n = 15) or WT copies for KMT2D (n = 15).

#### SUPPLEMENTARY TABLES

**Table S1: List of epigenetic modifiers and corresponding shRNAs used for RNAi screen.**

**Table S2: Pathway analysis of differential genes between KMT2D mutant versus wild type tumors in 6 different TCGA tumor types.**

**Table S3: List of primers and antibodies used in the study.**

**Figure S1**

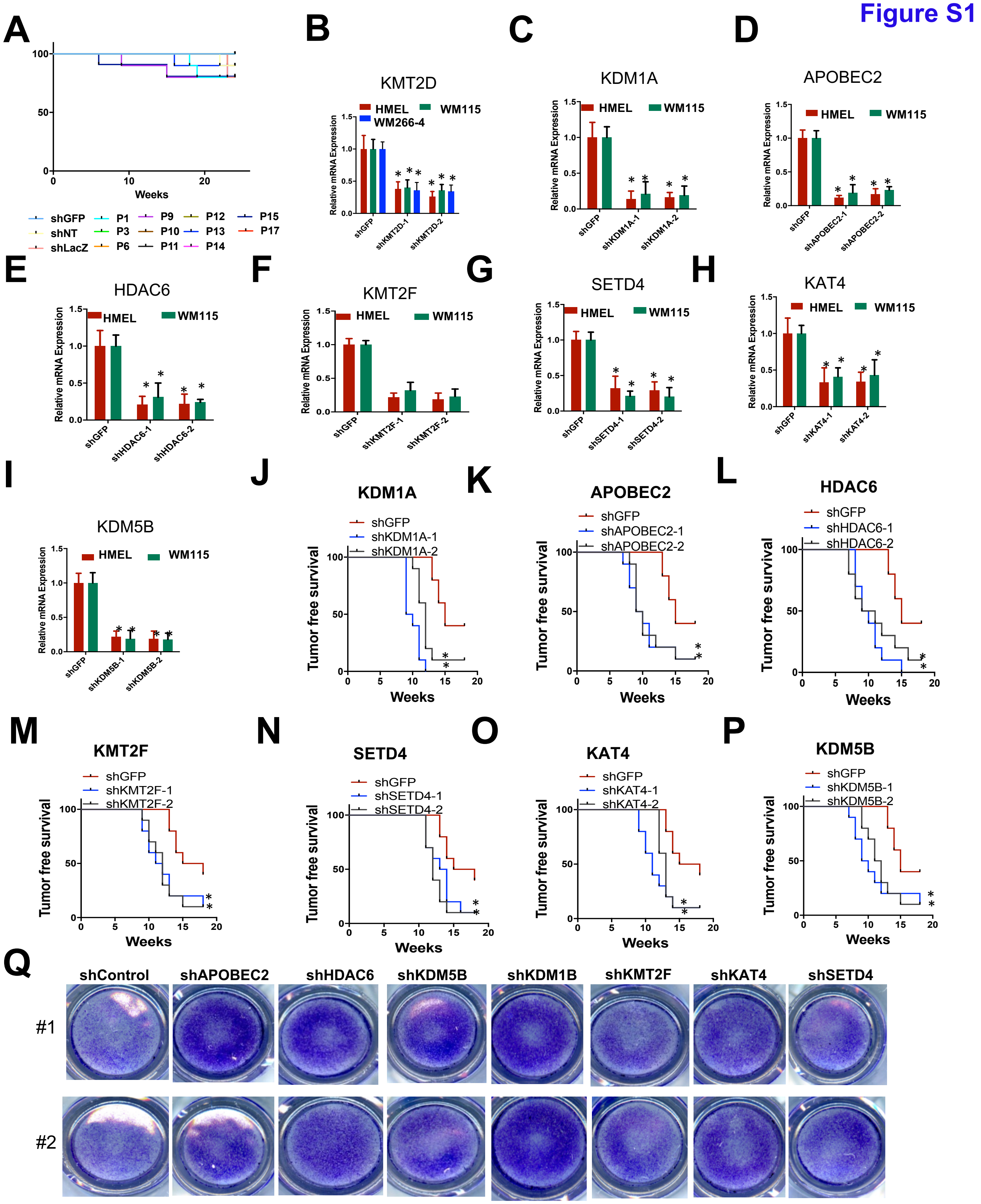


**Supplementary Figure S1: Validation of epigenetic regulators identified through RNAi screen identified 8 epigenetic regulators as potential tumor suppressors in melanoma.**

**a.** Kaplan-Meier curve showing tumor-free survival of mouse cohorts orthotopically injected with 1 million HMEL-BRAF$^{v600E}$ cells transfected with pooled shRNAs from primary screen. This figure shows data from pools that did not significantly accelerate tumorigenesis. Three negative control pools (shLuc, shGFP and shNT) are shown. Cells from each pool were injected in 10 mice each and tumor formation was monitored over 25 weeks.

**b-i.** Bar graph showing relative levels of indicated genes in the HMEL-BRAF$^{V600E}$ and WM115 cells harboring lentivirally integrated shRNAs for that gene or GFP. Y-axis represents fold change of the gene expression compared to 28s and normalized to control shRNA samples. Standard t-test $^{*}p < 0.05$, $^{**}p < 0.01$, $^{***}p < 0.001$

**j-p.** Kaplan-Meier curve showing tumor-free survival of mouse cohorts orthotopically injected with WM115 cells stably expressing shRNAs against KDM1A (**j**), APOBEC2 (**k**), HDAC6 (**l**), KMT2F (**m**), SETD4 (**n**), KAT4 (**o**) and KDM5B (**p**). Mantel-cox test $^{*}p < 0.05$, $^{**}p < 0.01$, and $^{***}p<0.001$.

**q.** Representative images for invaded cells from Boyden chamber assay of HMEL-BRAF$^{V600E}$ cells harboring lentivirally integrated shRNAs for GFP (control) or APOBEC2, HDAC6, KDM5B, KDM1A, SETD4, KMT2F or KAT4. #1 and #2 represent the duplicates for the experiment.

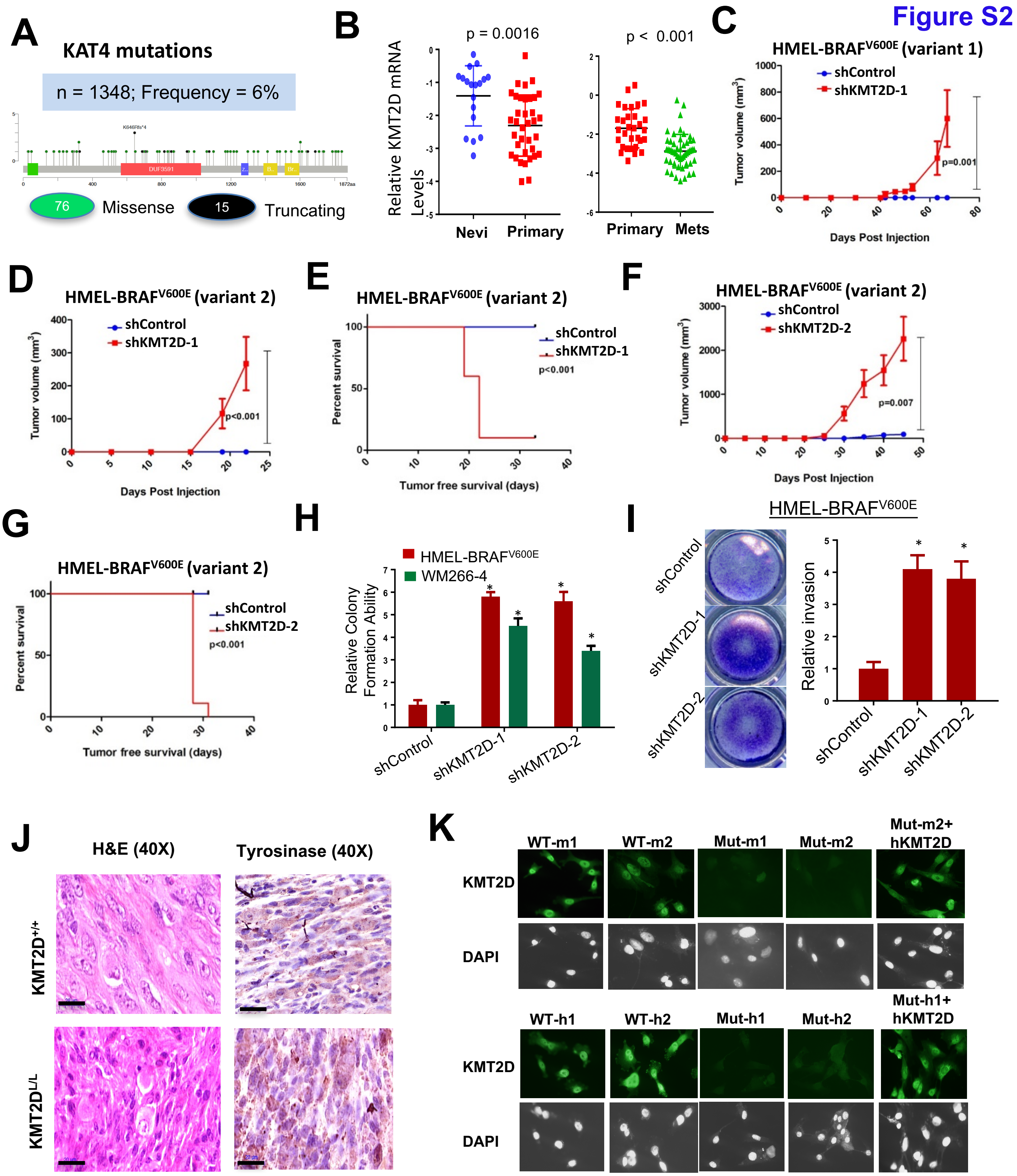


**Supplementary Figure S2: Validation of KMT2D as a tumor-suppressor in melanoma.**

**A.** Schematic of KAT4 protein showing missense mutations seen across all melanoma studies deposited in cBio portal. Green filled circles denote missense mutations whereas black filled circles represent truncating mutations Colored boxes within KAT4 schematic show different protein domains.

**B.** Dot plot showing relative KMT2D mRNA levels between nevi and primary melanomas (from Reference 21) (left panel) and between primary and metastatic melanomas (from Reference 22) (right panel).

**C, D, F.** Graph showing tumor volume of HMEL-BRAF$^{V600E}$ cells (clonal variant 1 in panel **C** and clonal variant 2 in panels **D** and **F**) harboring either control (shNT) or KMT2D shRNAs (shKMT2D-1 in panel **d** and KMT2D-2 in panel **f**).

**E, G.** Kaplan-Meier curve showing tumor-free survival of mouse cohorts orthotopically injected with HMEL-BRAF$^{V600E}$ cells (clonal variant 2) harboring either control (shNT) or KMT2D shRNAs (shKMT2D-1 in panel **e** and KMT2D-2 in panel **g**).

**H.** Relative soft agar colon formation ability of HMEL-BRAF$^{V600E}$ or WM266-4 cells harboring control or 2 different KMT2D shRNAs (shKMT2D-1 and shKMT2D-2). Unpaired t-test *p < 0.05, **p < 0.01, and ***p<0.001.

**I.** Representative images and quantification of invaded HMEL-BRAF$^{V600E}$ cells stably transfected with control or KMT2D shRNAs in Boyden Chamber assay. Unpaired t-test *p < 0.05, **p < 0.01, and ***p<0.001.

**J.** Images of H&E stained and Tyrosinase stained (standard Immunohistochemistry) (40X) melanoma tumors from *iBIP;KMT2D*$^{+/+}$ and *iBIP;KMT2D*$^{L/L}$ mice. Scale bar is 20μM.

**K.** Images of **i**mmunofluorescence for KMT2D in WT-m1, WT-m2, Mut-m1, Mut-m2, Mut-m2 + dox (hKMT2D), WT-h1, WT-h2, Mut-h1, Mut-h2, and Mut-h1 + dox (hKMT2D) cells for KMT2D. Nuclear staining is shown by DAPI staining (shown in grayscale).

**Figure S4**

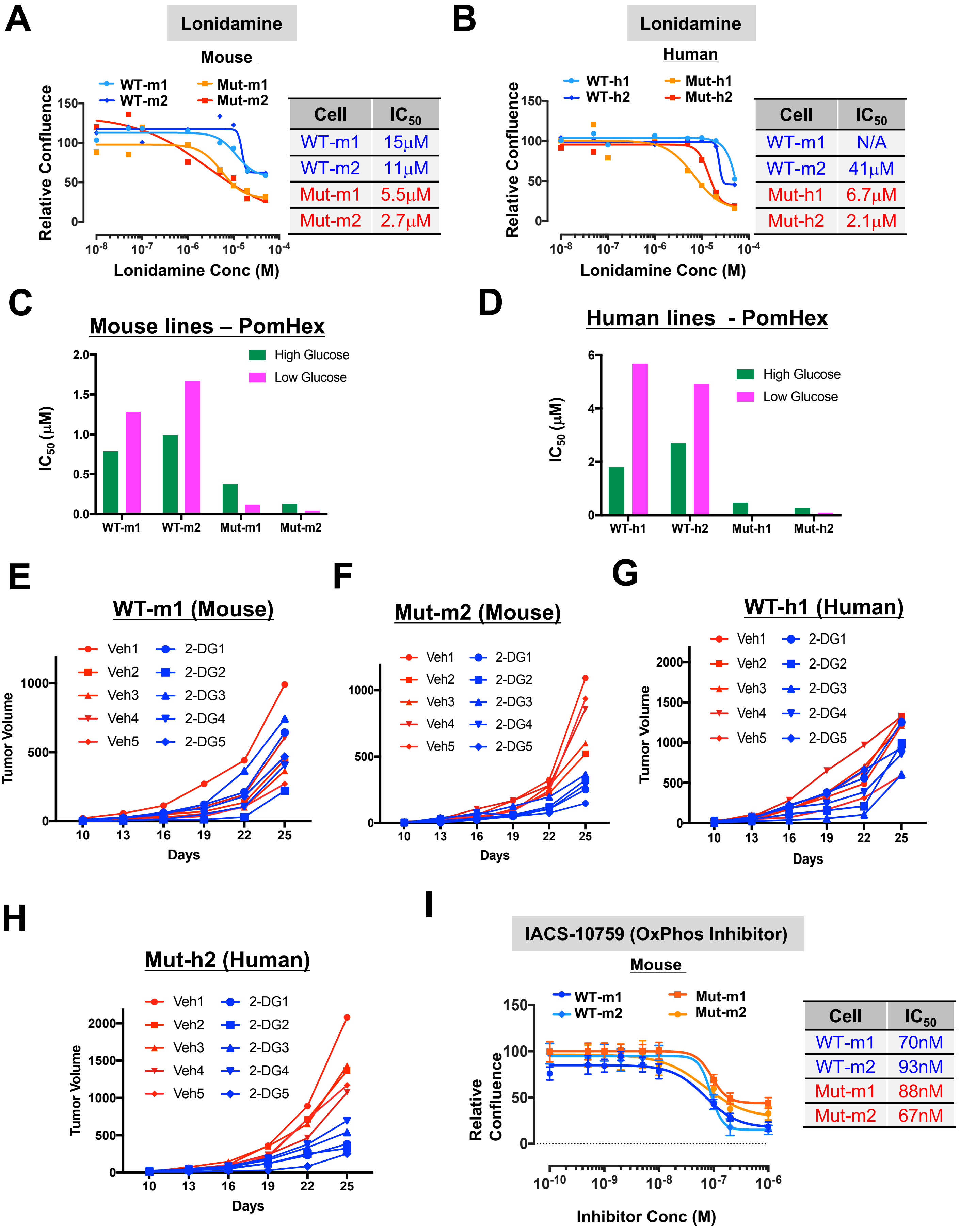


**Supplementary Figure S4: Inhibition of glycolysis preferentially impacts KMT2D mutant cells.**

**A-B.** Growth curves for KMT2D mutant and WT murine (**A**) and human (**B**) melanoma cells treated with varying concentrations of Lonidamine. Relative confluence at 96-hours post treatment are plotted and IC50 values are shown in the accompanying table.

**C-D.** Bar plot showing $IC_{50}$ values for Pomhex upon treatment of WT-m1, WT-m2, Mut-m1, Mut-m2 (**C**) or WT-h1, WT-h2, Mut-h1, Mut-h2 (**D**) cells in high glucose (4g/L) or low glucose (1g/L) DMEM media.

**E-H.** Line plot showing tumor volumes for individual mice (n = 5 per group) treated with 2-DG (500mg/kg) or vehicle every other day. Individual panels show data from mice injected with murine KMT2D WT (WT-m1) (**E**), murine KMT2D mut (Mut-m2) (**F**), human KMT2D WT (WT-h1) (**G**) and human KMT2D mutant (Mut-h1) (**H**) melanoma cells treated with vehicle or 2-DG.

**I.** Growth curves for KMT2D mutant and WT murine melanoma cells treated with varying concentrations of IACS-10759, an inhibitor of oxidative phosphorylation. Relative confluence at 96-hours post treatment are plotted and IC50 values are shown in the accompanying table.

### Figure S3

**A**

**GO DOWNREGULATED iBIP KMT2D MUT VS WT**

| GO Term | p-value |
|---|---|
| GO:0007155~cell adhesion | 1.7e-10 |
| GO:0009611~response to wounding | 1.3e-8 |
| GO:0042445~hormone metabolic process | 6.3e-7 |
| GO:0006954~inflammatory response | 3.5e-5 |
| GO:0000904~cell morphogenesis involved in differentiation | 1.7e-4 |

**B**

**HALLMARK ALL iBIP KMT2D MUT VS WT**

| NAME | NOM p-val | FDR q-val |
|---|---|---|
| HALLMARK_INTERFERON_GAMMA_RESPONSE | 0.0E+00 | 0.0E+00 |
| HALLMARK_INTERFERON_ALPHA_RESPONSE | 0.0E+00 | 0.0E+00 |
| HALLMARK_HYPOXIA | 0.0E+00 | 0.0E+00 |
| HALLMARK_MYOGENESIS | 0.0E+00 | 1.3E-02 |
| HALLMARK_TNFA_SIGNALING_VIA_NFKB | 0.0E+00 | 1.3E-02 |
| HALLMARK_IL6_JAK_STAT3_SIGNALING | 6.0E-03 | 1.3E-02 |
| HALLMARK_EPITHELIAL_MESENCHYMAL_TRANSITION | 4.0E-03 | 4.9E-02 |
| HALLMARK_GLYCOLYSIS | 8.2E-03 | 8.4E-02 |
| HALLMARK_WNT_BETA_CATENIN_SIGNALING | 4.6E-02 | 9.3E-02 |
| HALLMARK_UV_RESPONSE_DN | 4.5E-02 | 1.8E-01 |

**C**

**HALLMARK TCGA SKCM**

| Pathway | NOM p-val | FDR q-val |
|---|---|---|
| HALLMARK_ALLOGRAFT_REJECTION | 0.0E+00 | 0.0E+00 |
| HALLMARK_OXIDATIVE_PHOSPHORYLATION | 0.0E+00 | 0.0E+00 |
| HALLMARK_ESTROGEN_RESPONSE_LATE | 0.0E+00 | 0.0E+00 |
| HALLMARK_INFLAMMATORY_RESPONSE | 0.0E+00 | 0.0E+00 |
| HALLMARK_KRAS_SIGNALING_DN | 0.0E+00 | 2.5E-04 |
| HALLMARK_XENOBIOTIC_METABOLISM | 0.0E+00 | 4.0E-04 |
| HALLMARK_KRAS_SIGNALING_UP | 0.0E+00 | 8.6E-04 |
| HALLMARK_REACTIVE_OXIGEN_SPECIES_PATHWAY | 0.0E+00 | 1.7E-03 |
| HALLMARK_IL6_JAK_STAT3_SIGNALING | 0.0E+00 | 3.2E-03 |
| HALLMARK_COMPLEMENT | 0.0E+00 | 2.9E-03 |
| HALLMARK_ESTROGEN_RESPONSE_EARLY | 0.0E+00 | 3.0E-03 |
| HALLMARK_P53_PATHWAY | 0.0E+00 | 2.8E-03 |
| HALLMARK_APICAL_JUNCTION | 0.0E+00 | 3.5E-03 |
| HALLMARK_APICAL_SURFACE | 1.7E-03 | 3.7E-03 |
| HALLMARK_TNFA_SIGNALING_VIA_NFKB | 0.0E+00 | 3.7E-03 |
| HALLMARK_IL2_STAT5_SIGNALING | 0.0E+00 | 8.9E-03 |
| HALLMARK_COAGULATION | 3.0E-03 | 1.0E-02 |
| HALLMARK_FATTY_ACID_METABOLISM | 3.0E-03 | 1.6E-02 |
| HALLMARK_HYPOXIA | 0.0E+00 | 1.9E-02 |
| HALLMARK_INTERFERON_GAMMA_RESPONSE | 6.1E-03 | 2.9E-02 |
| HALLMARK_UV_RESPONSE_UP | 2.2E-02 | 5.2E-02 |
| HALLMARK_CHOLESTEROL_HOMEOSTASIS | 4.5E-02 | 5.3E-02 |
| HALLMARK_GLYCOLYSIS | 3.1E-02 | 9.2E-02 |
| HALLMARK_MYOGENESIS | 4.8E-02 | 9.9E-02 |

**D**

**HALLMARK_GLYCOLYSIS**

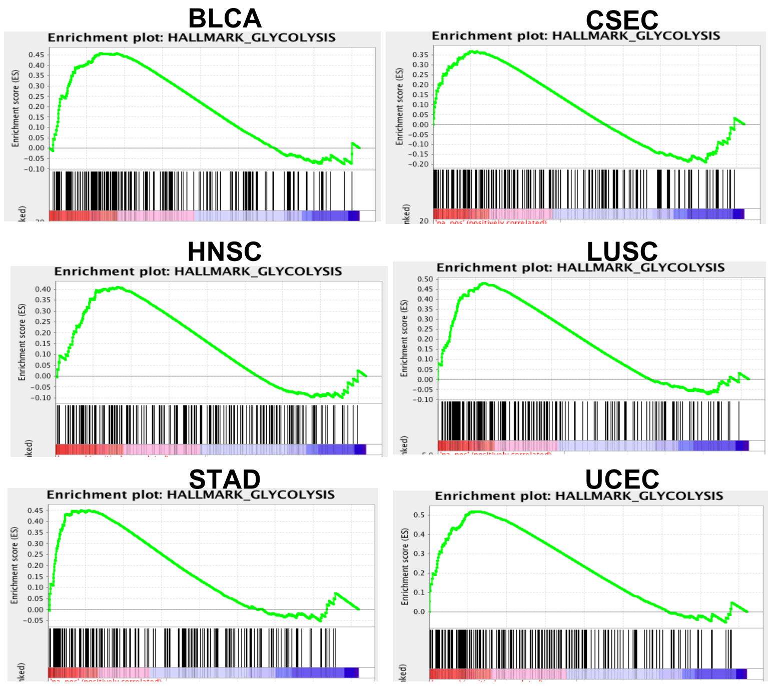


**E**

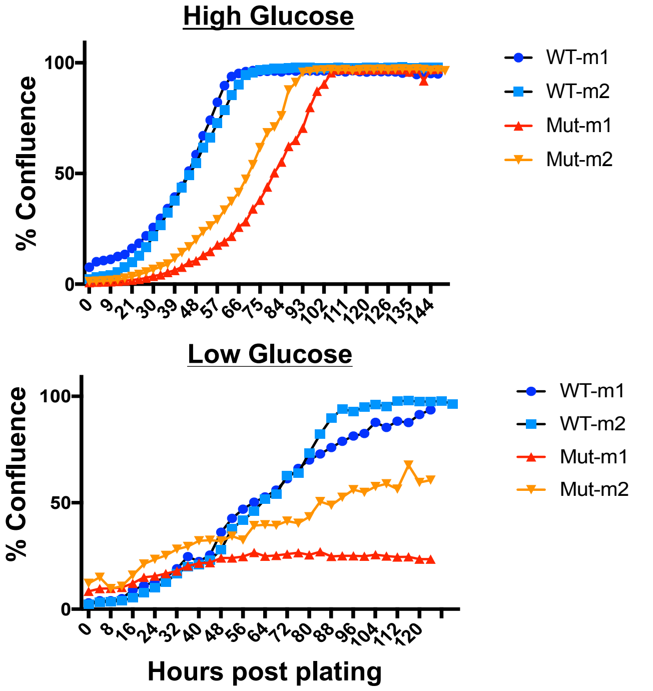


**F**

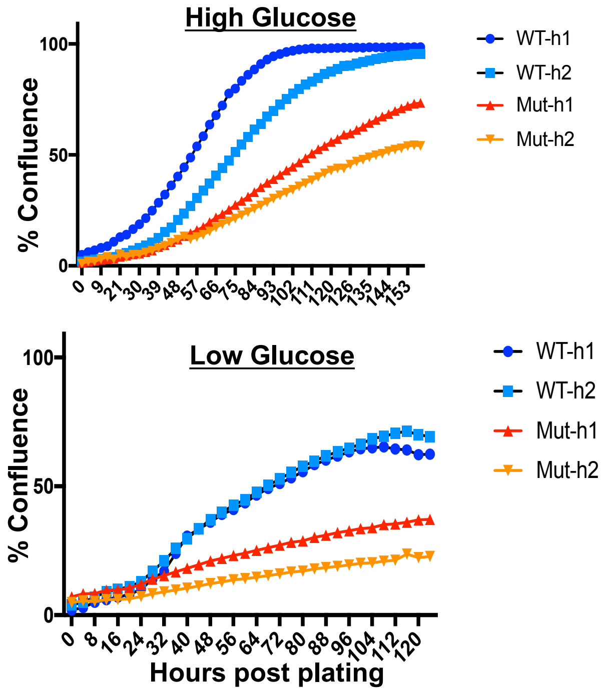


**Supplementary Figure S3: KMT2D mutant cells induce energy metabolism pathways and depend on glucose availability.**

**A-B.** Top 5 GO terms (**a**) and all significant HALLMARK terms for downregulated genes (FDR < 0.05, FC >2) (**b**) between KMT2D mutant murine cells and KMT2D wild type cells by total RNA-Seq analysis.

**C.** All significant HALLMARK pathway terms in differentially expressed genes between KMT2D mutant (carrying truncation, frameshift and post4700aa missense) and wild type human primary melanomas from the TCGA data.

**D.** Enrichment plots for HALLMARK glycolysis pathway in differentially expressed genes between between KMT2D mutant (carrying truncation, frameshift and post4700aa missense) and wild type human tumors from six different tumor types in TCGA data where n of mutant samples > 10.

**E-F.** Growth curves for WT-m1, WT-m2, Mut-m1 and Mut-m2 cells (**E**) and WT-h1, WT-h2, Mut-h1, Mut-h2 cells in DMEL high glucose (4g/L) or low glucose (1g/L) media. X-axis represent hours post plating and Y-axis shows percent confluence.

Figure S5

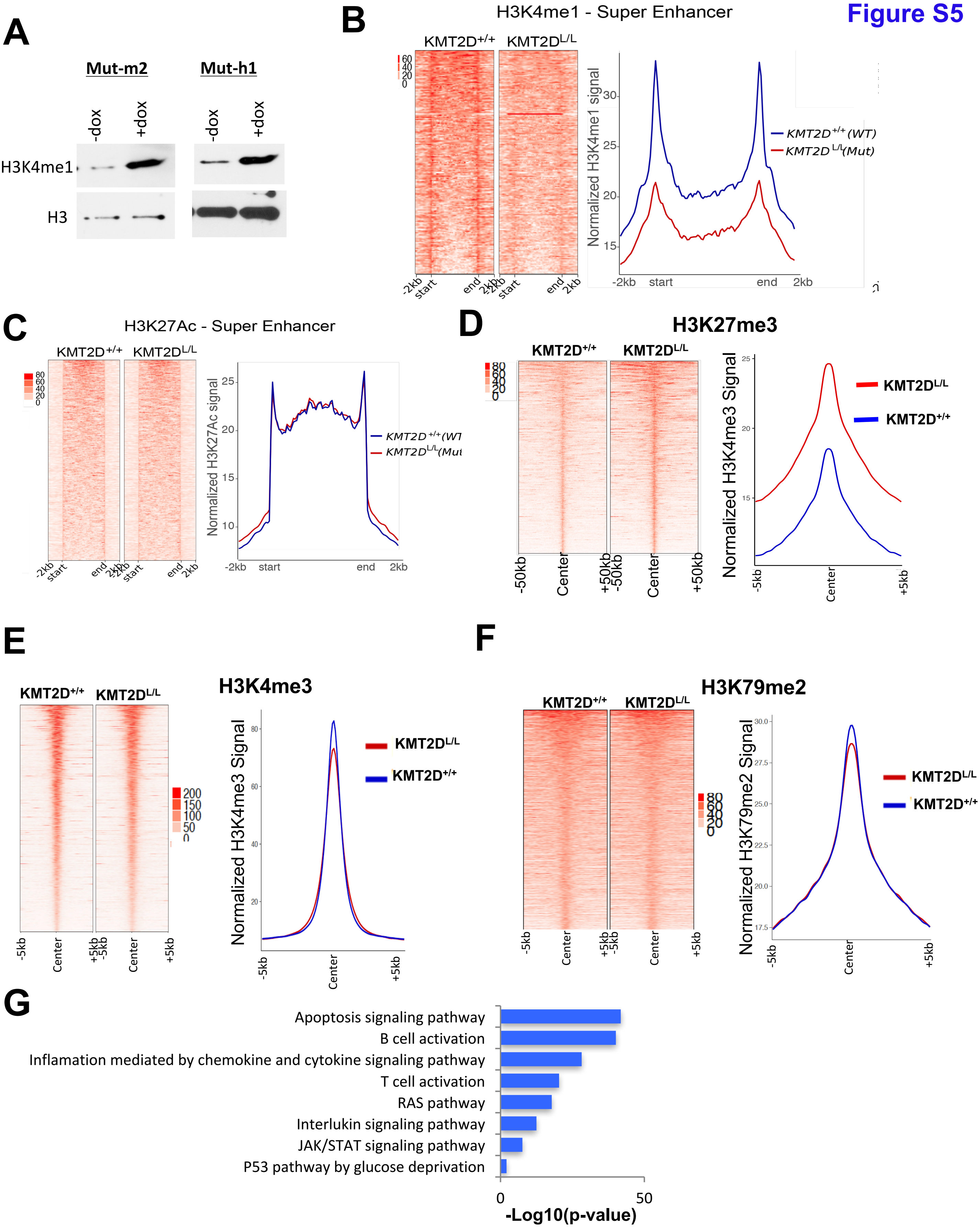


**Supplementary Figure S5: Levels of H3K4me3, H3K27me3 and H3K79me2 in KMT2D WT and mutant melanoma tumors.**

**A.** Western blot showing total H3K4me1 and H3 in KMT2D mutant (Mut-m2, Mut-h1) and rescue with hKMT2D (+dox) murine (left) and human (right) melanoma cells.

**B-C**. Heat map (left panels) and average intensity curves (right panels) for super-enhancer peaks based on H3K4me1 (**B**) and H3K27ac (**C**) ChIP-Seq data peak in *iBIP;KMT2D*$^{+/+}$ and *iBIP;KMT2D*$^{L/L}$ melanoma tumors.

**D-F.** Heat map (left panels) and average intensity curves (right panels) derived from the ChIP-Seq reads (RPKM) for H3K27me3 (**D**), H3K4me3 (**E**) and H3K79me2 (**F**) at enriched promoters in 10kb window centered on the middle of the peak in *iBIP;KMT2D*$^{+/+}$ and *iBIP;KMT2D*$^{L/L}$ melanoma tumors. Signals show enrichment in regions that enriched for these marks in respective type of tumor.

**G.** Pathway analysis for active enhancer (H3K4me1 and H3K27ac overlap) peaks specific to KMT2D mutant compared to KMT2D wild type tumors.

**Figure S6**

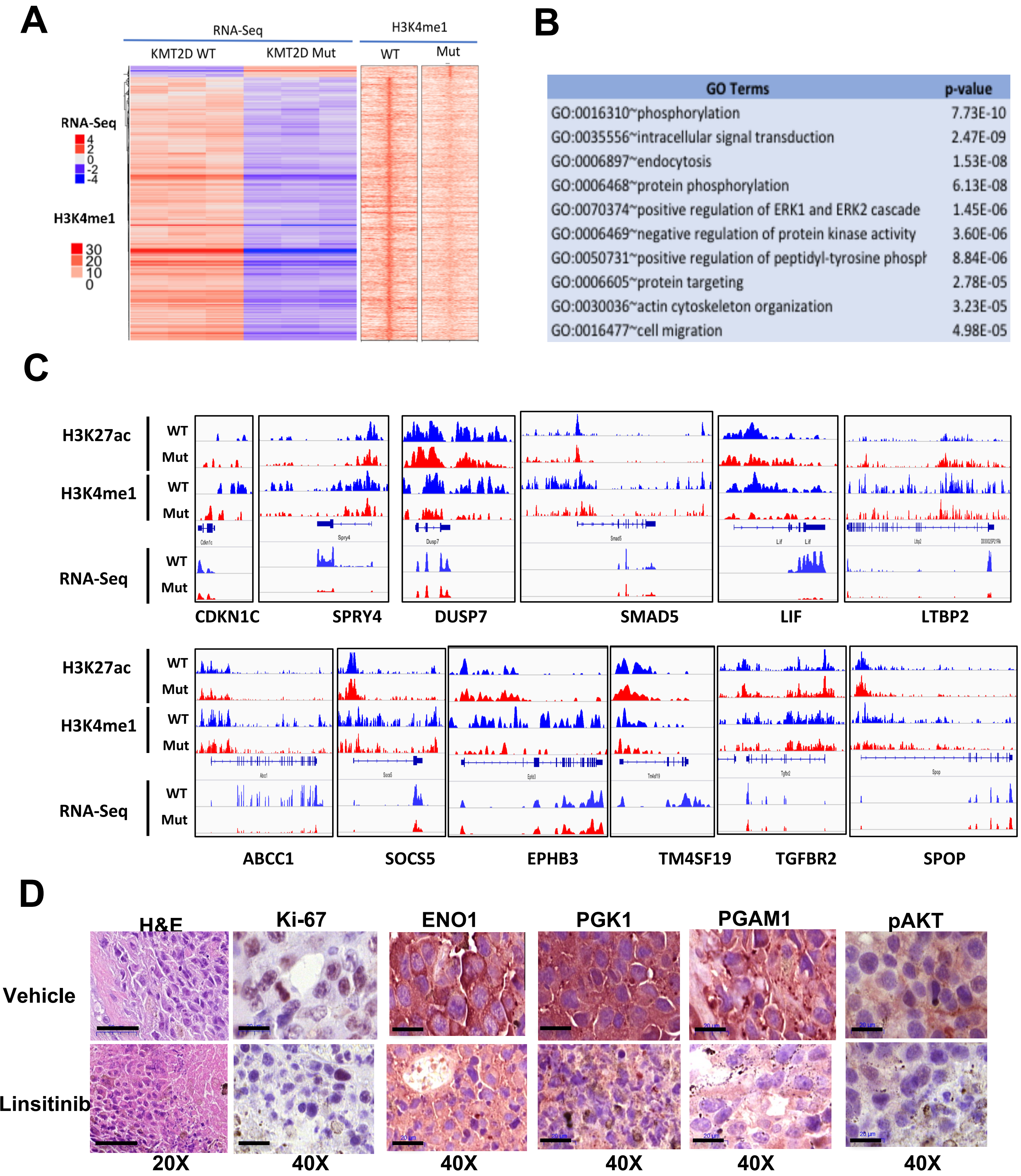


**Supplementary Figure S6: IGF signaling and aberrantly activates glycolysis in KMT2D mutant melanoma tumors.**

**A.** Heat maps showing differentially expressed genes and associated H3K4me1 signals in the vicinity as noted. R1, R2 and R3 refer to three replicates for WT-m1 and Mut-m2 cells.

**B.** All significant GO terms for genes which are differentially expressed and harbor loss of H3K4me1 mark at associated loci in KMT2D mutant melanoma cells compared to WT tumor cells.

**C.** Snapshots of IGV viewer for H3K4me1, H3K27ac and RNA-Seq signals on genomic loci surrounding various *bona fide* or putative tumor suppressor genes in KMT2D wild type or mutant tumors.

**D.** Images of H&E stained tumor or those stained for the product of three glycolysis genes (ENO1, PGK1 and PGAM1) or pAKT in xenograft tumors of Mut-m2 cells treated with vehicle or Linsitinib (20mg/kg). Magnification is shown below the images. Scale bar shows 20µM in all cases except H&E images where it represents 50µM.

**Figure S7**

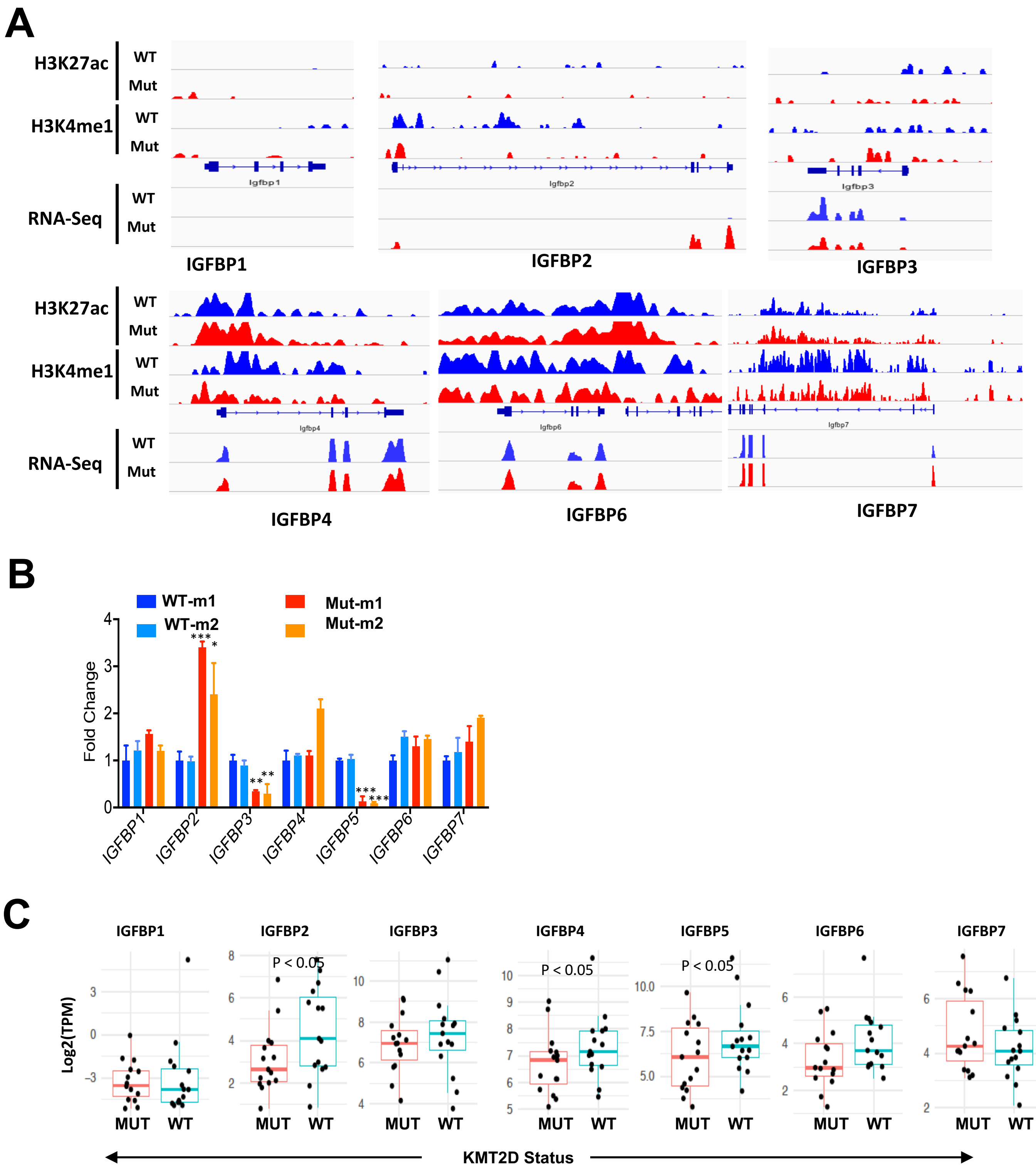


**Supplementary Figure S7: Loss of enhancer activity at IGFBP5 mediates activation of IGF signaling and aberrantly activates glycolysis in KMT2D mutant melanoma tumors.**

**A.** IGV snapshot showing RNA-seq, H3K27Ac, H3K4me1 ChIP-seq signal tracks for genomic locus associated with IGFBP proteins.

**B.** Bar graph showing relative expression levels of indicated IGFBP genes in murine KMT2D mutant (Mut-m1 and Mut-m2) and WT (WT-m1 and WT-m2) cells. Y-axis represents fold change of the gene expression compared to 28S and normalized to control shRNA samples. Standard t-test *p < 0.05, **p < 0.01, ***p < 0.001

**C.** Box plot showing expression of IGFBP genes in the melanoma TCGA samples that harbor functional mutations (nonsense, frameshift or post4700aa) (n = 15) or WT copies for KMT2D (n = 15).
